# Nuc3DMap delineates finer nucleosome-based three-dimensional chromatin architecture

**DOI:** 10.1101/2025.04.03.647017

**Authors:** Kun Fang, Lavanya Choppavarapu, Tianxiang Liu, Philip A. Knight, Victor X. Jin

**Affiliations:** Division of Biostatistics, Data Science Institute; MCW Cancer Center, Medical College of Wisconsin, Milwaukee, WI 53226, USA; Department of Mathematics and Statistics, University of Strathclyde, Glasgow G1 1XH, UK

**Author notes:** Correspondence: Victor X. Jin.

**Keywords:** nucleosome-based, three-dimensional, finer chromatin architecture

## Abstract

Currently, there lacks a method to delineate finer nucleosome-based 3D chromatin architecture. In this study, we develop a novel computational method, nucleosome-based 3D chromatin map (Nuc3DMap), to integrate MNase-seq, Hi-C and Micro-C data. Nuc3DMap implements an ensembled LOESS-sparse Knight-Ruiz algorithm to efficiently balance nucleosome-based contact matrix and adapts a bottom-up approach to identify nano-scale chromatin organization including nucleosome-based topological domain (NucD), boundary (NucB), gap (NucG) as well as nucleosome-based chromatin interaction loci (NucIL) and chromatin loop (NucL). We evaluate Nuc3DMap on publicly available data in H1 and GM12878 cells and identify the distinct subtypes of functional NucBs and NucLs, respectively, by integrating them with more than a hundred of multi-omics modalities data. Interestingly, functional NucBs exert distinct nucleosome organizational features and different insulation capability; and Pol2-related NucLs implicated Pol2 as a noncanonical regulator in mediating gene looping and cell type specificity. Furthermore, our work systematically demonstrates the relationship between interaction intensity and transcriptional activity, unveiling an activation-like behavior of chromatin looping. Overall, Nuc3DMap represents a conceptual advancement in chromatin conformation analysis, offering a refined view of genome organization from the fundamental nucleosome level to higher-order chromatin structures, as well as serves as a foundational framework for advancing computational methods’ development in dissecting nano-scale 3D genome organization, enabling new discoveries and fostering innovation in chromatin organization modeling.

## Main

Chromosomal conformation capture (3C)-based genome-wide techniques such as Hi-C have enabled us to survey the three-dimensional (3D) organization of the genome within a living cell at a level of detail that was previously impossible, providing a comprehensive view of how DNA is folded and interacts with itself in a cellular environment^1^. The dynamics of genome organization has been linked to cellular function in many biological processes^2^. It has become clear that the 3D chromatin architecture can be distinguished at three levels, including A and B compartments corresponding to active and repressive chromatin segments, respectively^3^, topologically associating domains (TADs) and structural loops extruded by the structural proteins CTCF and cohesin^4–8^, and local chromatin domains or chromatin loops such as enhancer–promoter loops (EPLs) and promoter–promoter loops (PPLs) often mediated by transcriptional regulators (TranRs) and epigenetic regulators (EpiRs)^9–11^. Although Hi-C can identify all these chromatin structures, it suffers from the relatively low resolution due to the restriction enzymes that give a bias on fragmentation of chromatin. Thus, the coverage of Hi-C is not comprehensive and uniform^12^. Micro-C, a recent renovated technique, is capable of effectively resolving ultra-fine 3D chromatin architecture at single nucleosomes^13^. Despite of this enhanced resolution, Micro-C limits in the detection of relatively shorter range of chromatin interactions. Therefore, an integration of Hi-C and Micro-C together will provide a holistic view of genome organization.

Nucleosomes and nucleosome-free regions form the primary architecture of single chromatin fibers, serving as a fundamental basis for genome function^14^. Nucleosome positioning acts as a barrier to promoter escape by Pol2^15^, and the +1 nucleosome is tightly bound to the preinitiation complex (PIC)-mediator complex, underscoring its organizational role in regulating the transcription process^16^. Two tetra-nucleosome folding motifs have been identified in the yeast genome^17^, and nucleosomes are arranged in heterogeneous clutches in mammalian cells^18^, further supporting the existence of higher-order micro-architectures at the nucleosome level. Although Micro-C utilizes MNase to digest the genome, the conventional fixed-bin contact matrix does not fully exploit the nucleosome-specific information embedded in this technique. Our previous study demonstrated that nucleosome-based binarization improves the identification of genomic features and their associated biological functions^19^. While a few tools, such as iNucs^20^, can build contact matrices using nucleosome locations, they exclude nucleosome-free regions and are therefore unable to capture potential interactions between transcription factors (TFs) in nucleosome-free regions and adjacent nucleosomes. Thus, it is critical to incorporate nucleosomes and their derived intervals from MNase-seq into contact matrices which facilitate a bottom-up approach to dissect chromatin architectures.

Over the past one decade and a half, numerous computational methods have been developed to delineate 3D chromatin architecture at each of the three levels^21^. For example, many tools, such as Cooltools^22^, Juice-Box^23^, HOMER^24^ and Fan-C^25^, derived, defined and implemented compartments by the sign of the first principal component analysis (PCA) on transformed Hi-C matrices. More than two dozen programs have been developed to predict TADs^5–6,26–29^. Several tools, including FitHiC^30^, HiSIF^31^ and MUSTACHE^32^, have been implemented to identify chromatin interactions or loops. There are some pioneering works in exploring computational methods for integrating Hi-C data with various multi-omics data^33–36^. However, there lacks a method to delineate finer nucleosome-based 3D chromatin architecture by integrating MNase-seq, Hi-C and Micro-C data.

In this study, we develop a novel computational method, <u>nuc</u>leosome-based <u>3D</u> chromatin <u>map</u> (Nuc3DMap), *i*.*e*., a nucleosome-based 3D contact map by computationally integrating MNase-seq, Hi-C and Micro-C data. Nuc3DMap comprises five modules, including NucPrep, NucLoad, NucMerge, NucDom and NucIL. Nuc3DMap implements an ensembled LOESS-sparse Knight-Ruiz algorithm to efficiently balance nucleosome-based contact matrix and adapts a bottom-up approach to identify key chromatin organization including nucleosome-based topological domain (NucD), boundary (NucB), gap (NucG) as well as nucleosome-based chromatin interaction loci (NucIL) and chromatin loop (NucL). We first evaluate Nuc3DMap on publicly available data in H1 cells, and further test and apply it on the data in GM12878 cells. We further integrate the NucB and NucL with more than a hundred of multi-omics modalities data to identify the distinct subtypes of functional NucBs and NucLs, respectively. We finally demonstrate that Nuc3DMap is capable of delineating nucleosome-based 3D chromatin architecture in a cell-type-specific manner.

## Results

### Overview of Nuc3DMap

Nuc3DMap employed a nucleosome-based genome segmentation approach as well as an ensembled LOESS-sparse Knight-Ruiz scaling method to construct a nucleosome-based 3D contact map by effectively integrating MNase-seq, Hi-C and Micro-C data (**Fig. 1A-B**). The Nuc3DMap further facilitated the identification of chromatin organization features, e.g., TAD and interaction loci at nucleosome resolution with several refined methods (**Fig. 1C-D**). The input of Nuc3DMap includes MNase-seq used for detecting nucleosome locations^37–39^, and Hi-C and Micro-C used for identifying genome-wide chromatin interaction^3,6,12–13^. Specifically, the Nuc3DMap comprises five modules, NucPrep, NucLoad, NucMerge, NucDom and NucIL (**Fig. 1**). Briefly, NucPrep processes MNase-seq, Hi-C, and Micro-C raw reads to determine nucleosome positions via the iNPS^40^ and to identify ligation junctions with Pairstool^41^ (**Methods**). NucLoad generates nucleosome-base contact matrices by nucleosomal binarization for Hi-C and Micro-C data, respectively, where such binarization can preserve the linear landscape information of the genome for further constructing chromatin contact matrices. NucMerge normalizes these matrices by using digestion sites-based LOESS normalization into balanced contact matrices and then integrates two matrices into a merged contact matrix devoid of systemic biases with the sparse Knight-Ruiz method. NucDom applies a sliding diamond-shaped window on the merged contact matrix to detect NucD, NucB, and NucG with significant features through curve-fitting and Mann-Whitney tests. NucIL applies a scale-invariant feature transformation (SIFT) with extended quantile detection on a renormalized grouping contact matrix to identify NucIL and NucL.

**Fig. 1.**
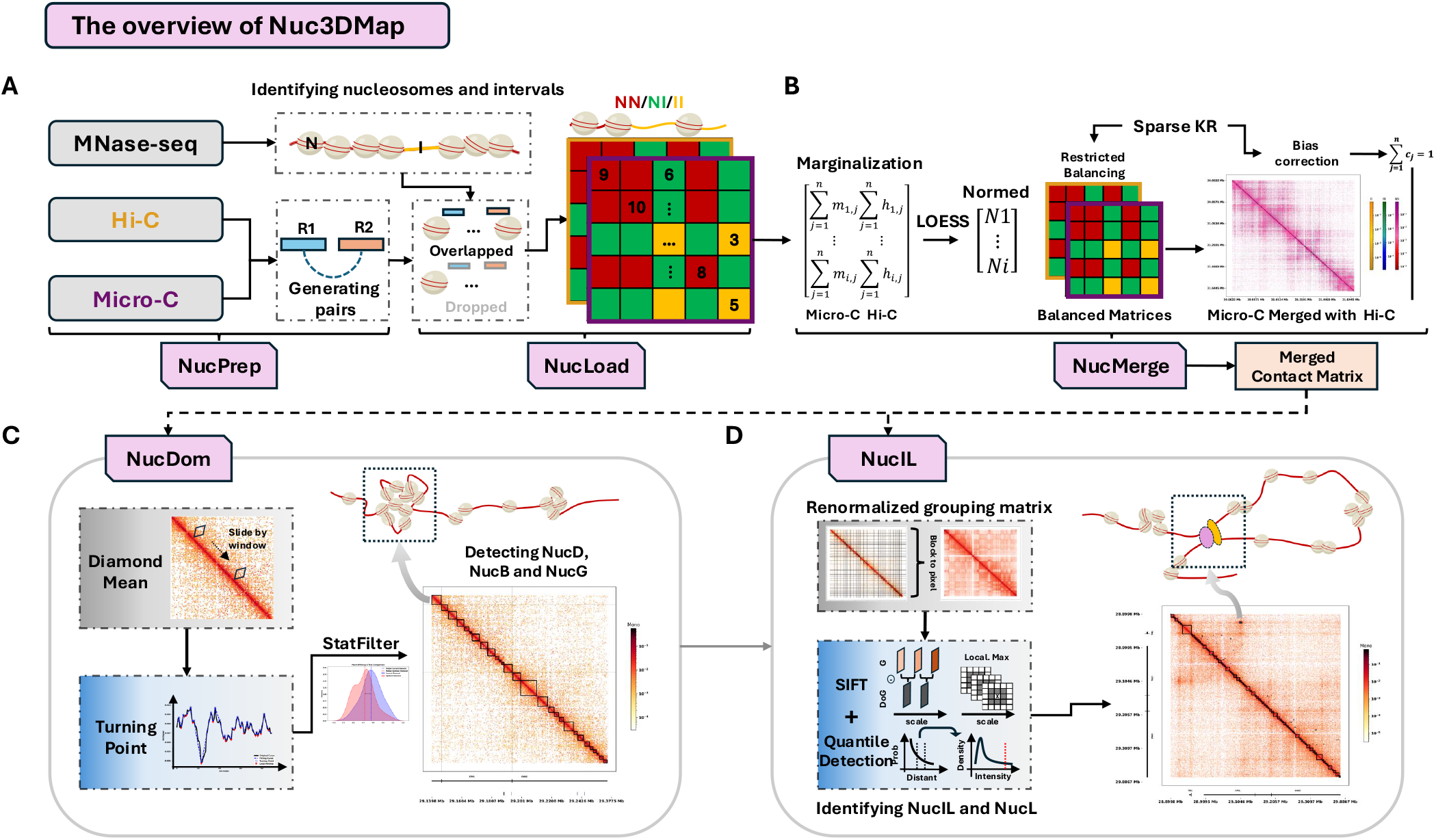
The overview of Nuc3DMap comprising five modules: NucPrep, NucLoad, NucMerge, NucDom and NucIL. **A**. NucPrep preprocessing uniquely mapped sequencing reads of MNase-seq, Hi-C and Micro-C data for determining nucleosome (N) positions from MNase-seq data and inferring intervals (I) information from Hi-C and Micro-C. Upon preprocessing, NucLoad constructs a nucleosome-based contact matrix based on the output from NucPrep by using nested containment (NCLS) list to efficiently quantify the number of pairs associated with each pixel (N or I) in the matrix. Red color represents N-N interaction; Green color represents N-I interaction; and yellow color represents I-I interaction. **B**. NucMerge generating a merged nucleosome-based contact matrix by normalizing and integrating Hi-C and Micro-C nucleosome-based contact matrices. **C**. NucDom identifying NucD, NucB and NucG on the merged nucleosome-based contact matrix from NucMerge. **D**. NucIL using a tri-step process to detect NucIL.

### Efficient and accurate construction of nucleosome-based contact matrices

We built and trained Nuc3DMap on MNase-seq, Hi-C and Micro-C data in H1 cells (**Extended Data Table 1**). Upon preprocessing the data with NucPrep (**Extended Data Fig. 1A**), we identified 11.89 million genome-wide nucleosomes from MNase-seq data and processed approximately 2.25 and 1.97 billion interacting pairs for Hi-C and Micro-C data, respectively (**Extended Data Fig. 1B**). NucLoad first segmented the whole genome into one-dimensional (1D) nucleosome-based bins according to the location of detected nucleosomes (**Fig. 2A**). There are two types of bins: nucleosome (N) bins and interval (I) bins, where N-bins were derived from the extended core locations of nucleosomes identified by NucPrep by applying a more stringent filtering process to filter out highly fuzzy nucleosomes (**Methods**). After extending and filtering, we obtained a total of 11.58 million nucleosome bins (**Fig. 2B**), with a median size of 160 bp (**Fig. 2C** – top-left), an averaging positioning score of 5.62 (**Fig. 2C** – bottom-left, **Methods**) and an averaging spacing of 238.44 bp (**Extended Data Fig. 1C**). The I-bins were further divided into short segments with relatively comparable size (a median size of 180 bp) to N-bins (**Fig. 2C** – top-right) and further classified into four categories based on the intensity of nucleosomes at the regions and the length of the regions (**Extended Data Fig. 1D, Methods**), including nucleosome-free regions (NFR), nucleosome low-intensity regions (NLIR), nucleosome-depleted gaps (NDG) and desert regions (DRs) (**Fig. 2C** – bottom-right, **Extended Data Fig. 1E**). We observed that NFRs were accounted for more than 50% of the I-bins, followed by DRs and NLIRs, while NDGs were the smallest proportion (**Extended Data Fig. 1F**).

**Fig. 2.**
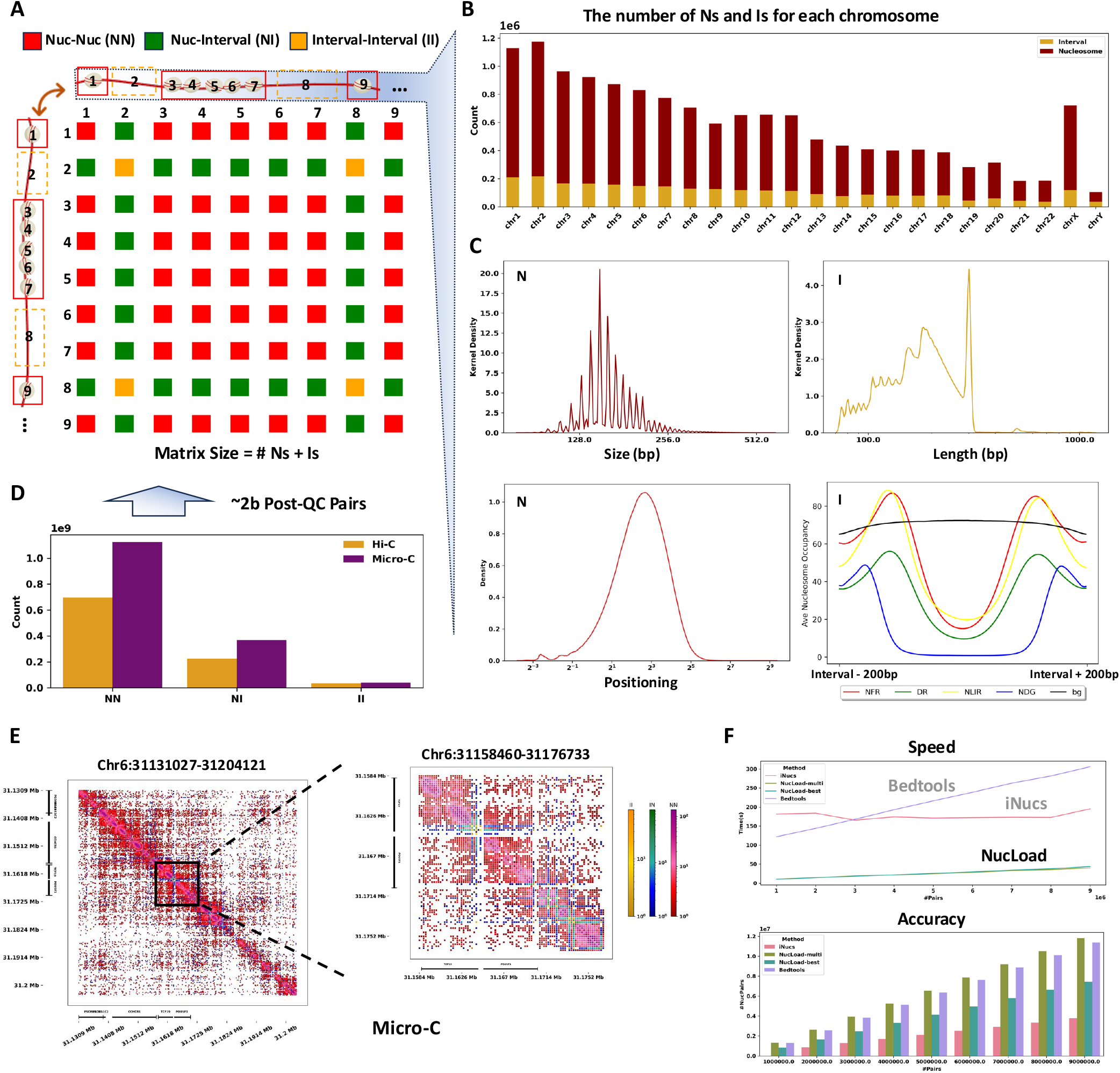
Efficient and accurate construction of nucleosome-based contact matrices by NucLoad. **A**. A schematic illustration of the construction of a nucleosome-based contact matrix. NucLoad segments the genome into two types of bins: N and I, with I was defined as the region between two consecutive nucleosomes that meet specific criteria (see **Methods** in detail). In the resulting 2D contact map, there are three distinct types of interactions: nucleosome-nucleosome (NN), nucleosome-interval (NI), and interval-interval (II) interactions. These are represented in the schematic by red, green, and yellow squares, respectively. **B**. The bar plot quantifying the number of nucleosome and interval bins across each chromosome. **C**. The top-left panel illustrating the size distribution of N-bins; the top-right panel displays the size distribution of I-bins; the bottom-left panel depicts the positioning scores of N-bins; and the bottom-right panel compares four types of I-bins characterized by low MNase signal with 200 bp extension, relative to random bins. **D**. The bar graph illustrating the total number of interaction pairs categorized into three types: nucleosome-nucleosome (NN), nucleosome-interval (NI), and interval-interval (II) interactions. Hi-C data is represented in orange while Micro-C data is depicted in purple. **E**. Visualization of the H1 Micro-C nucleosome-based contact matrix across genomic regions Chr6:31131027-31204121 and Chr6:31158460-31176733. Three major colors, yellow, green, and red, represent the types of interaction, II, NI, and NN, respectively. **F**. A comparison of performance among NucLoad, iNucs, and Bedtools in constructing nucleosome-based contact matrices. The upper panel line plot displays that NucLoad builds the contact matrix more than 10x faster than iNucs and Bedtools do. The bottom bar plot shows that NucLoad’s best-match mode achieves approximately 2 times higher accuracy compared to iNucs, while NucLoad’s multi-mode exhibits a performance comparable to Bedtools’ intersection method. These evaluations are based on artificially constructed datasets ranging from ~1 to 9 million pairs (see **Methods** in detail).

After processing the N-bins and I-bins, NucLoad built and populated a nucleosome-based contact matrix by incorporating the ordered combination of the N-bins and I-bins (referred to as ‘bins’ for the convenience, **Fig. 2A**) and chromatin interaction pairs (**Fig. 2D**). A big challenge in this process was to assign interaction pairs efficiently and accurately to pixels within the nucleosome-based contact matrix, where a pixel represents an entry in the matrix associated with the genomic coordinates of interaction pairs. Current fixed-bin based methods^23,42,43^, *i*.*e*., counting the 5’ end of reads into fixed bins, were not suitable for discontinuously variable-sizes bins which might lead to contact pairs loss and inaccuracies. NucLoad thus employed an interval intersection-based method, cored with the nested containment list algorithm^44^, to precisely assign the billions of interaction pairs into nucleosome-based bins. This approach was able to efficiently identify pairs with at least a 1bp overlap with the bins. Furthermore, NucLoad implemented a multi-step overlapping method and found the best match for nucleosome assignment (**Extended Data Fig. 1**, and **Methods**). Finally, NucLoad completed the nucleosome-based contact matrix by assigning all bin-associated pairs to the appropriate pixels, as illustrated in **Fig. 2E**. Intriguingly, we observed that there are some blocks formed by N-I pixels (**Fig. 2E**) suggesting that intervals themselves might conditionally act as domain boundaries^45^. Additionally, we conducted a comparison of the performance among NucLoad, Bedtools, and iNucs using a series of synthetic datasets with varying numbers of interaction pairs (**Methods**). Our analysis showed that NucLoad achieved approximately ten times faster in speed and nearly two times greater in accuracy compared to iNucs under various parameter settings (**Fig. 2F** and **Extended Data Fig. 1G**). All in all, NucLoad efficiently and accurately constructs nucleosome-based contact matrices that preserve genome information, including critical interval regions and further offers a much faster and more accurate performance compared to existing tools.

### Normalization and integration of nucleosome-based contact matrices

Hi-C and Micro-C have been shown to exhibit distinct interaction frequency patterns that reflect differences in resolution and experimental design^12,13,46–48^. We also found that the chromatin interaction pairs of Micro-C displayed an increased interaction frequency within a shorter range of genomic proximity while the interaction pairs of Hi-C were more across larger genomic distances (**Fig. 3A**). Further examination of the distribution of genomic distances for NN, NI, and II interactions revealed varied interaction frequency patterns (**Fig. 3B**). For example, interaction frequencies of common pixels observed in both Hi-C and Micro-C spanned from ~100 bp to ~1 Mb. Interaction frequencies of Hi-C-specific pixels had two respective peaks at ~100 kb and ~10 Mb while interaction frequencies of Micro-C-specific pixels displayed two distinct peaks at ~1 kb and ~100 kb. Interestingly, a middle range of interaction frequencies at ~100kb were captured in both Hi-C-specific and Micro-C-specific pixels. To integrate Hi-C and Micro-C data efficiently and accurately into a unified nucleosome-based contact matrix and to ensure that both data types are at a comparable scale, NucMerge employed a three-step normalization process: digestion site (DS)-Anchor identification (**Fig. 3C**), DS-Anchor LOESS Scaling, and sparse Knight-Ruiz (sKR) Balancing (**Fig. 3D**). We observed a higher correlation between Hi-C and Micro-C data across genomic regions containing one DS and more (**Fig. 3C**), suggesting that the presence and density of DSs could be used for the alignment of interaction data between the two data types. We defined regions, *i*.*e*., nucleosome-based N and I bins, with at least one DS as normalization anchors, termed DS-Anchors. These DS-Anchors in each chromosome were scaled using LOESS to adjust discrepancies between Hi-C and Micro-C data, aiming to re-level the fitting line close to zero (**Methods, Fig. 3D** – left, **Extended Data Fig. 2A**). For regions without DS-Anchors, we applied a linear interpolation of the scaling factors derived from the DS-Anchors to achieve scaling across the genome. NucMerge further utilized the scaled marginal values to perform the sparse Knight-Ruiz (sKR) Balancing to address the inherently sparse nature of our nucleosome-based contact matrix. This sKR Balancing consists of three key steps: connected components reordering, blocks balancing with sKR, and traceback genomic order (**Methods, Fig. 3D** – right). We tested the performance of the sKR Balancing and confirmed it is superior to the Row-Column Sum (RAS, akin to Vanilla-Coverage) method in terms of convergence and to the original KR algorithm in terms of speed (**Fig. 3E, Extended Data Fig. 2B**,**C**). NucMerge was able to scale the marginal values and sequentially normalize the contact matrices of both Hi-C and Micro-C data (**Fig. 3F, Extended Data Fig. 2D**,**E**). NucMerge finally incorporated the contact frequency of Hi-C-specific pixels into the normalized Micro-C contact matrix (**Fig. 3G**), where the distinctive features of both Hi-C and Micro-C data were effectively retained as shown in **Fig. 3H**. A merged contact matrix with genomic bias implicitly removed with sKR balancing was showed in **Fig. 3I**. In summary, NucMerge ensures a robust merged nucleosome-based contact matrix with combined strengths of both Micro-C and Hi-C techniques.

**Fig. 3.**
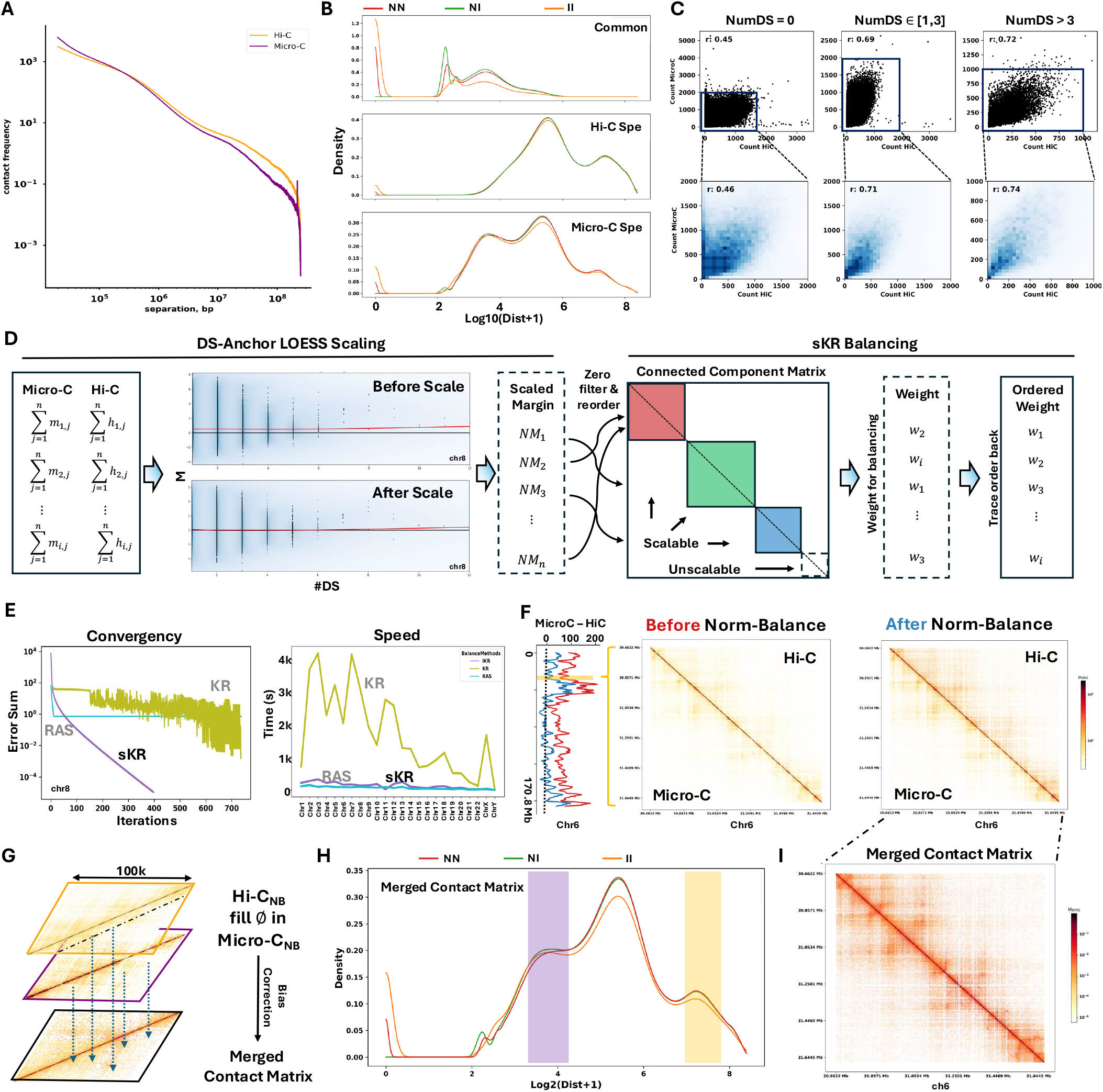
Integration of a nucleosome-based contact matrix by NucMerge. **A**. Interaction frequency between Hi-C and Micro-C data being plotted on a Log10 scale against genomic distance for H1 cells. **B**. The distribution of genomic distances for NN, NI, and II pixel types in nucleosome-based contact matrices being displayed with color-coded in red, green, and yellow, respectively. The common pixels, Hi-C-specific and Micro-C-specific pixels were arrangement from the top to the bottom in the plot, demonstrating unique and shared interaction patterns. **C**. Scatterplots in the upper panel depicting the correlations of marginal values for different genomic segmentations categorized by the number of DSs (DS=0, DS ∈ [1,3], DS>3) between Hi-C and Micro-C data. The lower panel shows density plots of these correlations on anchors identified across different DS (DS-Anchor). **D**. Two normalization steps in NucMerge starting with DS-Anchor based LOESS scaling of the marginal values for both Hi-C and Micro-C data, followed by an application of sparse Knight-Ruiz (sKR) normalization, which includes connected component reordering and non-bistochastic KR balancing to enhance matrix equilibration. **E**. A comparison of performance of the sKR method against KR and RAS methods in terms of convergence and computational efficiency, showing sKR is superior in convergence in fewer iterations and shorter time, with RAS failing to converge. **F**. The left line plot depicting the re-leveling of marginal values to zero via NucMerge normalization across genomic regions. The right panel visually demonstrates the effect of this normalization, showcasing enhanced data uniformity and balance post-normalization. **G**. A visualizing plot of the merging process in NucMerge. In the merged matrix, non-zero pixels from Micro-C retained their original values, ensuring that fine-scale interactions were preserved. Null pixels with middle and long-distance, where no data is available from Micro-C data, were supplemented with values from Hi-C data to provide a comprehensive view of chromosomal interactions at both short and long genomic distances. **H**. The distribution of genomic distances for NN, NI, and II pixel types within the merged nucleosome-based contact matrix. The pixel types are color-coded as red for NN, green for NI, and yellow for II. This visualization demonstrates the merged contact matrix is capable of preserving interactions across both short-range and long-distance interactions at the same matrix. **I**. The visualization of a merged nucleosome-based contact matrix. The color indicates the intensity of interaction frequency.

### Identification of nucleosome-based topological domain and boundary

TADs are self-interacting genomic regions and their boundaries divide the genome into distinct regulatory territories^4,5,49,50^. However, the determination of TADs and its boundaries is largely dependent on the resolution of the contact matrix and sensitive to the algorithms and their corresponded parameters^51,52^. Leveraging the nucleosome-partitioned information preserved in the nucleosome-based contact matrix, we further implemented the NucDom module to identify NucD, NucB, and NucG (**Methods**). To achieve the best performance of NucDom, we first optimized the only parameter, window size (Winsize) through testing a series of Winsize ranging from 3 to 20 pixels. We observed that a variable size of the NucDs and a variable number of NucDs at different Winsize with a peak at 10 (**Extended Data Fig. 3A**). We then devised a scoring method for each Winsize by summing ranks assigned in a descending order based on the number of CTCF peaks (**Extended Data Fig. 3B**) and the median discrepancy between intra- and inter-contact frequencies at each NucB (**Extended Data Fig. 3C**). We thus determined a Winsize of 10 as the optimal and default parameter (**Extended Data Fig. 3D**). In addition, we also assessed the robustness of sequencing depth of NucDom (**Extended Data Fig. 4A**). Our robust analysis revealed that NucDom integrity was primarily affected by variations in Micro-C data, with a lesser impact observed from Hi-C data adjustments (**Extended Data Fig. 4B-E**). Further analysis of CTCF peak overlap with NucB across conditions revealed a significant increase between Micro-C at 25% and 50% sequencing depth (**Extended Data Fig. 4F-G**). This suggests that achieving optimal performance with NucDom requires more than 1.5 billion post-quality control reads for Micro-C.

In H1 cells, we identified ~559k NucDs, ~512k NucBs and ~48k NucGs (**Extended Data Table 2**), each with varied distribution in size (**Fig. 4A**) and in interaction frequencies (**Extended Data Fig. 3E**). Notably, the median size of a NucB is 325 bp, approximately the length of two nucleosomes, suggesting that NucDom can identify nucleosome-based boundary (**Extended Data Fig. 3F**). We further classified NucB based on many publicly available genome-wide data of TranRs and EpiRs, including five 3D chromatin structure-related transcription factors (STF)^6,7,53–57^, CTCF, RAD21, SMC3, YY1 and ZNF143, 26 histone marks (HIS), 36 TF, 3 chromatin remodelers (CR), 12 histone transferases (HISase), R-Loop (RLoop), 5-DNA methylation (5mC), 5-hydroxyl-DNA methylation (5hmC) and nascent RNA (nasRNA) of H1 cells (**Extended Data Table 1, Extended Data Fig. 3G**). We surprisingly found that only 12.8% NucBs were associated with one STF or two more combinatorial (c)STFs, where the top four cSTFs were CTCF-RAD21, CTCF-RAD21-SMC3, CTCF-RAD21-SMC3-ZNF143 and CTCF-RAD21-SMC3-YY1-ZNF143 (**Fig. 4B**). We further examined the NucBs with 13 types of combinatorial TranRs and EpiRs, including active histone marks (A-HIS), repressive marks (R-HIS), poised histone marks (P-HIS), elongation histone marks (E-HIS), Pol2, TFSS/TFNS, HISase, CR, STF, RLoop, 5mC, 5hmC and nasRNA, and measured the correlation between each of two types (**Fig. 4C**). Intriguingly, we observed higher correlations between 5hmC and A-HIS and between nasRNA and Pol2, whereas neither RLoop nor 5mC had any correlations with other TranRs. For these correlation coefficient score larger than or equal to 0.5, we derived 12 combinatorial patterns, including A~STF, P~STF, A~5hmC, P~5hmC, A~STF~5hmC, P~STF~5hmC, A~STF~5hmC~nasRNA, P~STF~5hmC~nasRNA, A~Pol2~STF~5hmC, P~Pol2~STF~5hmC, A~Pol2~STF~5hmC~nasRNA, and P~Pol2~STF~5hmC~nasRNA (**Methods**). We then defined 25 types of functional NucBs with distinct genomic features and combinatorial patterns (**Fig. 4D, Extended Data Fig. 5A** and **Extended Data Table 3**). For example, E-HIS NucBs were enriched at gene body, P~STF NucBs were almost evenly distributed on distal, promoter and gene body while a majority of A~Pol2~STF~5hmC~nasRNA NucBs were located at promoter. Interestingly, we observed RLoop NucBs were the highest among all of NucBs, indicating its potential role as a chromatin insulator.

**Fig. 4.**
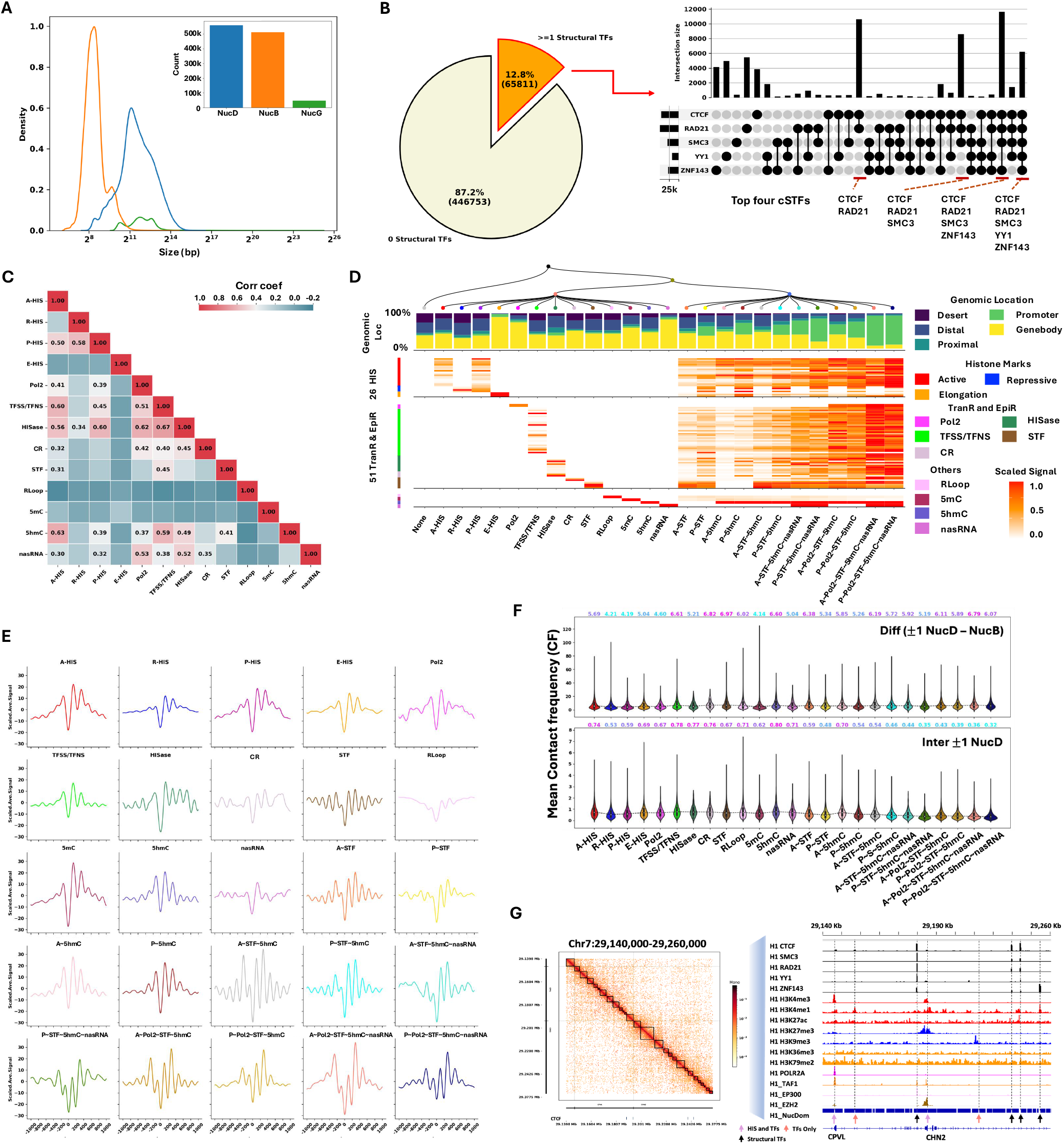
Identification of NucDs and NucBs in H1 cells. **A**. Line and bar plots depicting the size distribution and a total number of NucD, NucB, and NucG, respectively. **B**. Left: Pie chart showing the proportion of NucBs associated with STFs. Right: UpSet plot displaying combinations of STFs within STF-related NucBs. **C**. The heatmap representing the correlation metrics across 13 ‘single’ type of NucBs. **D**. The multi-feature plot showing a detailed genomic characteristic of 25 types of distinct functional NucBs, including genomic location (desert, distal, proximal, promoter and gene body), associated HIS, TF, CR, HISase, RLoop, 5mC, 5hmC, and nasRNA. **E**. The MNase density plot representing the nucleosome organization adjacent to each type of NucB. **F**. Top: Violin plots comparing contact frequency discrepancies between different NucB types and flanking NucDs, annotated with mean values and color intensity indicating higher values. Bottom: Violin plots of interacted contact frequencies between NucDs flanking various NucBs **G**. IGV screenshots of identified NucD and NucB within a specific region on chromosome 7, illustrating precise TFs captured by NucBs.

Next, we quantitatively assessed the organization features of nucleosomes flanking around NucB including nucleosome phasing, spacing, and positioning and revealed diverse organizational patterns (**Fig. 4E, Extended Data Fig. 5B-D, Extended Data Table 4**). We reaffirmed well-phased nucleosome arrays adjacent to STF^58^ and poorly phased arrays next to R-Loop and nascent RNA regions. Interestingly, we identified two primary modes of nucleosome organization among different types of functional NucB: Symmetric where nucleosome organization on both sides of the boundary is similar, and Asymmetric, where nucleosome organization on right side is more well-phased than left side. STF-related NucBs predominantly showed symmetric patterns, suggesting a directional influence of different NucB types. Conversely, other types exhibited asymmetric patterns, indicating that within a nucleosome domain (NucD), the beginning nucleosomes were more phased than those at the end. Further, we quantified the insulation capability by measuring the contact frequency difference between NucBs and the flanking NucDs, as well as the contact frequency (CF) between two adjacent NucDs. Intriguingly, combinatorial NucB generally demonstrated better insulation capability characterized by higher CF discrepancies between NucBs and NucDs, and lower CF among adjacent NucDs (**Fig. 4F**), indicating the insulation capability might be linearly correlated with the number of TFs or complex on the boundary. Additionally, we recapitulated that the number of nucleosomes within Pol2-associated NucDs were less than the ones of non-Pol2-associated NucDs^18^ (**Extended Data Fig. 5E**). Notably, we also found precise alignment of CTCF and other TFs at NucBs (**Fig. 4G**), suggesting NucDom effectively captured potential binding patterns of CTCF and TFs at nucleosome-based resolution. A comparison with conventional boundaries identified using a fixed bin method from the 4DN database^1^ revealed our NucBs exhibited more centralized alignment of CTCF than those identified by a fixed bin method (**Extended Data Fig. 5F**). Overall, NucDom can identify nucleosome-based topological domains and boundaries with an unprecedented resolution, where distinct subtypes of functional NucBs are associated with distinct nucleosome organizational features and different insulation capability.

### Identification of nucleosome-based interaction loci

We next implemented the NucIL module to identify the NucIL. We first transformed the sparse nucleosome-based contact matrix into RG-Map by utilizing a renormalization grouping concept^59^ (**Extended Data Fig. 6A**). This transformation effectively reduced the matrix size by approximately 100 times, facilitating a more efficient examination of nucleosome interactions without losing any essential genomic information. We further employed a dual-step method with optimized parameters to capture NucIL on the RG-Map (**Methods, Extended Data Fig. 6B-F**). The effectiveness of these parameters was confirmed by an Aggregated Peaks Analysis (APA) score of 21.5 as shown in **Extended Data Fig. 6G**. Additionally, the analysis revealed that NucIL integrity was influenced by variations in both Micro-C and Hi-C data, with a synergistic effect leading to fewer NucILs at lower data resolution for both methods (**Extended Data Fig. 4H**), further supporting the notion that the combination of Hi-C and Micro-C benefits the comprehensive detection of NucIL.

In total, we identified ~386k NucILs for H1 cells (**Fig. 5A, Extended Data Fig. 7A** and **Extended Data Table 5**). There are six types of NucIL, including NucD-NucD, NucD-NucB, NucD-NucG, NucB-NucB, NucB-NucG and NucG-NucG. Of all NucILs, NucD-NucB and NucB-NucB types accounted for the majority of interactions at ~130k and ~232k, respectively, suggesting that NucB might play a dominant role in folding 3D chromatin architecture. We next examined the length of each of six different NucIL types, defined as the distance between two loci, as well as the loci size of each type (**Fig. 5B** and **Extended Data Fig. 7B**). We observed a bi-modal distribution of the length for NucB-NucB type with a median more than twice longer (~106 kb) than that of NucD-NucD and NucD-NucB types (~47 kb), suggesting that NucD is more engaged in proximity interactions. Additionally, the median loci size for NucB-NucB is around 250 bp, approaching mono-nucleosome resolution. We further performed an autoencoder (AE) and Gaussian mixture model (GMM) clustering (**Extended Data Fig. 7C-E**) on NucIL with 14 TranRs and EpiRs, including five STFs, Pol2, and TAF1, three active marks (H3K4me3, H3K4me1, H3K27ac), two elongation marks (H3K36me3, H3K79me2), and two repressive marks (H3K27me3, H3K9me3). We obtained 14 clusters with distinct combinatorial patterns of 14 TranRs and EpiRs (**Fig. 5C**). Of 14 clusters, clusters 0, 2, 5, 10, 12, 13 were exclusively associated with histone marks, clusters 1, 4 were specific to TranRs, and clusters 3, 6, 8, 9, 11 exhibited a combination of both histone marks and TranRs. Cluster 7 showed little association with any TranRs. We also observed diverse interactions between STF and Pol2. CTCF, RAD21, and SMC3 tended to form clusters independent from Pol2, such as clusters 1, 4, 6, 11, while YY1 and ZNF143 were preferentially clustered with Pol2 and active histone marks, such as H3K4me3 and H3K27ac in cluster 9. Interestingly, cluster 3 was associated with CTCF, RAD21, SMC3, Pol2, TAF1, and H3K4me3, while cluster 8 exhibited Pol2 activity without the involvement of STF. These 14 clusters varied in number of interaction types, with more than half of each cluster comprising NucB-NucB interactions (**Fig. 5D**). To assess a similarity of nucleosome organization (NucOrg) between NucIL clusters, we calculated the scaled Fréchet distance (**Methods, Extended Data Fig. 7F**) and found that all clusters had a similarity score above 0.8 (**Fig. 5E**), suggesting topologically linked genomic regions share similar nucleosome organization features.

**Fig. 5.**
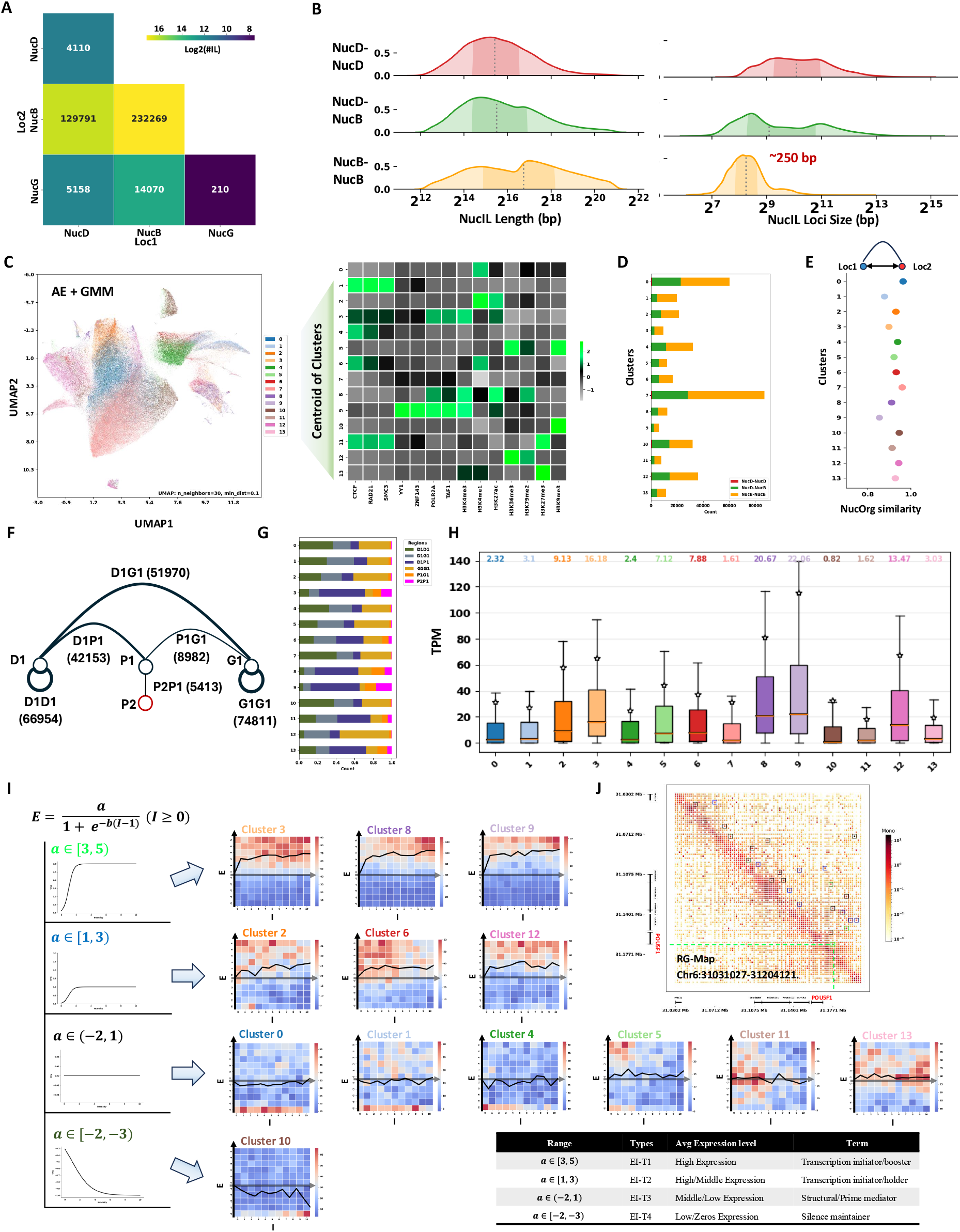
Identification and clustering of NucILs in H1 cells. **A**. The heatmap illustrating the numbers of each of six NucIL types in H1 cells: NucD-NucD, NucD-NucB, NucB-NucB, NucD-NucG, NucB-NucG, and NucG-NucG. **B**. The left KDE-plot showing the distribution of the length of each of three NucIL types: NucD-NucD, NucD-NucB, and NucB-NucB, where the length is the distance between two interaction loci. The right plot showed the distribution of the size of interaction loci of each of these three types, where the median size of NucB-NucB approaches 250 bp. **C**. The UMAP (left) visualizing 14 clusters of NucILs classified by using an AutoEncoder (AE) for dimensional reduction and a Gaussian Mixture Model (GMM) for clustering based on the selected histone marks and transcription regulators at their interaction loci. The heatmap (right) displays the centroids of each cluster, highlighting distinct combinatorial patterns of histone modifications and transcription factors. **D**. The bar plot illustrating the composition of each of three NucIL types, NucD-NucD, NucD-NucB, and NucB-NucB in each of 14 clusters. **E**. The dot plot demonstrating the NucOrg similarity measured by Fréchet distance between interaction loci from different NucIL clusters. **F**. The numbers of NucLs, a gene-centric type of NucILs categorized based on the genomic location of their interaction loci within protein-coding genes, distinguishing among D, P, and G, including six types of NucL, D1D1, G1G1, D1P1, D1G1, P1G1, and P2P1. **G**. The bar plot showing the count of each of six types of NucL in each of 14 NucIL clusters. The count is normalized to 1 for each of clusters. **H**. The box plot presenting the distribution of normalized gene expression in TPM for P-centric NucL, D1P1, P1G1 and P2P1 in each of 14 NucIL clusters. The average values were showed on the top. **I**. A non-linear sigmoid function being modeled on the interaction-expression matrix, resulted in four classes of relationships between chromatin interaction and gene expression (EI-T1, EI-T2, EI-T3 and EI-T4). Each of 14 clusters was able to fall in one of four classes. **J**. A visualization on the RG-Map highlighting identified NucLs with different rectangles indicating various types of chromatin interaction-gene expression relationships in color: lime for EI-T1, blue for EI-T2, black for EI-T3, and dark green for EI-T4.

We further categorized NucIL into NucL based on the genomic location of each locus related to an annotated gene: distal (D), proximal/promoter (P), and gene body (genebody, G). We obtained six types of NucL, including 74,811 G1G1, 68,954 D1D1, 51,970 D1G1, 42,153 D1P1, 8,982 P1G1 and 5,413 P1P2 loops, respectively (**Fig. 5F**). Interestingly, P1P2 loops have been reported to cluster multiple genes and provide topologically transcriptional coordination^60,61^. We also found that a diverse distribution of each of six types of NucL in each of 14 clusters (**Fig. 5G**). For example, D1P1 loops were composed of more than half of the NucLs in the Pol2-related clusters, 3,8,9, re-affirming an enhancer-promoter regulation mechanism^62^. Additionally, we observed an increasing proportion of P2P1 loops within these Pol2-related clusters while D1D1, D1G1, and G1G1 loops were predominantly STF-only clusters, 1 and 4. These findings prompted us to examine the relationship between P-centric NucL (D1P1, P1G1 and P2P1) and their associated gene’s expression within each of 14 clusters. As expected, Pol2-related clusters were associated with higher TPM (transcripts per million) values compared to non-Pol2-related clusters (**Fig. 5G**). Interestingly, genes associated with Pol2-related NucLs exhibited higher TPM values than those with Pol2 bound at their promoter but lacking NucLs (**Extended Data Fig. 7H**). These data indicate a critical role of NucL in regulating gene expression.

We further quantitatively modeled the relationship between P-centric NucL intensity and gene expression. We first scaled the TPM and NucL intensity using zTPM and Box-Cox transformations. Interestingly, both features exhibited a similar bimodal distribution after scaling (**Extended Data Fig. 7H**), suggesting a positive correlation between the two. We then constructed an expression-intensity (EI) matrix fitting a mean-value line based on the matrix (**Methods, Fig. 5I**) and modeled it using a modified sigmoid activation function with two parameters: *a*, the height of the plateau, and *b*, the slope of the linear part. Intriguingly, we identified four distinct classes associated with different P-centric NucL clusters. Specifically, we observed a high plateau curve for Pol2-related clusters (3, 8, 9) with *a* value within the range [3, 5), termed as EI-T1; a moderately high plateau for clusters associated with active and elongation histone marks (2, 6, 12) with *a* value within the range [1, 3), termed as EI-T2; a flat plateau for clusters associated with STF-only and STF-H3K27me3 (0, 1, 4, 5, 11, 13) with *a* value within the range (−2, 1), termed as EI-T3; and a negative plateau for the H3K9me3-related cluster (10) with *a* value within the range [−2, −3], termed as EI-T4. Our quantitative modeling suggests that specific TranRs and EpiRs might regulate the gene expression within each of four classes, EI-T1, EI-T2, EI-T3, and EI-T4. We thus defined each of four specific sets of TranRs and EpiRs as transcription initiator or booster, transcription holder, structural/prime mediator, and silence maintainer, respectively (**Fig. 5I**). We speculate that the class EI-T1 might be associated with cell type specificity. For example, POU5F1, a key embryonic stem cell gene, has an EI-T1 NucL (**Fig. 5J**). In summary, the NucIL module is able to identify NucIL and NucL, where the NucL can be further classified by various transcriptional and epigenetic regulators into distinct classes of functional NucLs in regulating specific subsets of gene transcription.

### Delineation of cell-type-specific nucleosome-based 3D chromatin architecture

3D Chromatin architecture often shows cell type specificity through interplaying with gene regulation program specifically to each cell type^55,63–67^. To test the efficiency of Nuc3DMap in a different cell type and delineate cell-type-specific 3D chromatin architecture, we applied it to a lymphoblastoid cell line, GM12878 which has been extensively studied in genomic research^68,69^. We identified ~330k NucD, ~290k NucB, and ~40k NucG in GM12878 cells (**Extended Data Table 3**), with a slightly different length distribution with more variation observed within NucD compared to H1 cells (**Fig. 6A**). Like H1 cells, only 15.6% of NucB in GM12878 were associated with STF with a low CTCF-RAD21 combination compared to H1 cells (**Extended Data Fig. 8A, B**). We then classified NucB with the same set of TranRs and EpiRs publicly available in GM12878 cells except 5hmC and RLoop data due to their unavailability, and identified 11 single type, A-HIS, R-HIS, P-HIS, E-HIS, Pol2, TFSS/TNFS, HISase, CR, STF, 5mC and nasRNA and 8 combination types A~STF, P~STF, A~STF~nasRNA, P~STF~nasRNA, A~Pol2~STF, A~Pol2~STF, P~Pol2~STF, A~Pol2~STF~nasRNA and A~Pol2~STF~nasRNA (**Extended Data Fig. 8C**,**D**). For NucB, we identified ~420k specific to H1 cells, ~94k shared between H1 and GM12878 cells, and ~220k specific to GM12878 cells (**Fig. 6B**). Interestingly, most shared NucBs related to STF, A-HIS and R-HIS were the primary contributors to H1-specific while 5mC and A~Pol2~STF~nasRNA were predominant in GM12878-specific NucB (**Fig. 6B**). When visually examining two typical cell-type-specific genes, POU5F1 for H1 cells and RUNX1 for GM12878 cells, we observed that a NucB along with Pol2, H3K4me3, and H3K27ac signals was present at the transcription start site (TSS) of POU5F1 in H1 cells and RUNX1 in GM12878 cells, respectively. However, NucB was absent neither at the POU5F1 site in GM12878 cells or at the RUNX1 site in H1 cells (**Fig. 6C**). This observation suggests that Nuc3DMap can effectively delineate cell-type-specific chromatin architecture at the level of nucleosomal domains and boundaries. We further identified ~318k NucILs in GM12878 cells (**Extended Data Table 6**) and a majority of them are the NucD-NucB and NucB-NucB types, each with ~144k and ~137k, respectively (**Fig. 6D**). We found a similar distribution for these two types as in H1 cells in terms of length and loci size (**Extended Data Fig. 8E**) and identified 16 clusters with distinct transcriptional and epigenomic patterns (**Extended Data Fig. 8F, G**). We also observed four expression-intensity patterns, EI-T1 to EI-T4, for NucILs in GM12878 cells as did in H1 cells (**Extended Data Fig. 8H**). These findings suggest that Nuc3DMap can effectively detect NucILs across different cell types.

**Fig. 6.**
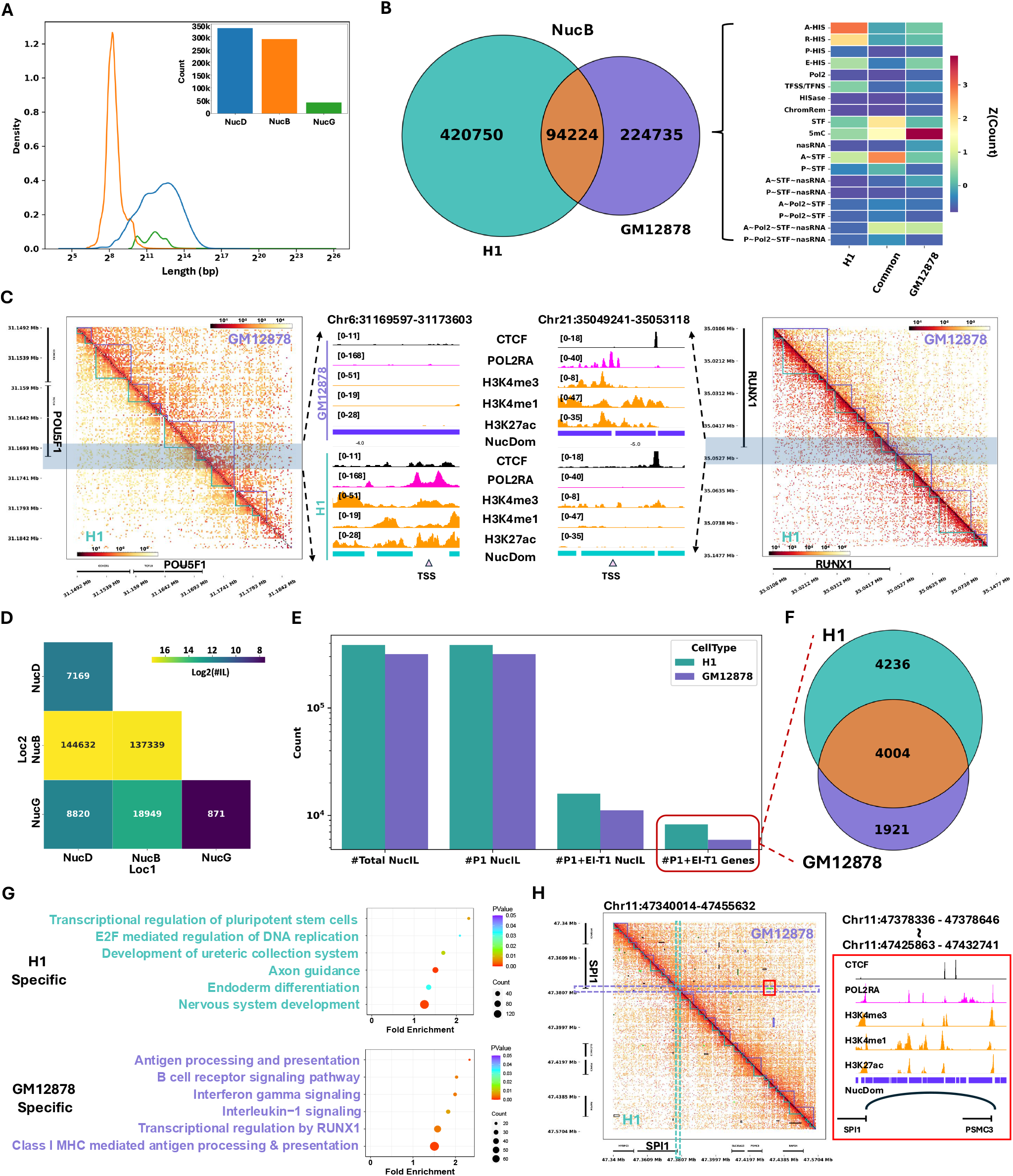
Cell type-specific nucleosome-based chromatin architecture in H1 and GM12878 cells, respectively. **A**. Line and bar plots depicting the length distribution and the total number of NucDs, NucBs, and NucGs in GM12878 cells, respectively. **B**. The Venn diagram (left) displaying the overlapping number of NucBs between H1 and GM12878 cells. The heatmap (right) visualizes the normalized count of H1-specific, Common and GM12878-specific NucBs in each of distinct functional NucB. **C**. The nucleosome-based contact maps and integrated genomics viewer (IGV) tracks for a H1-specific gene, POU5F1, and a GM12878-specific gene, RUNX1 in H1 and GM12878 cells, respectively. NucD regions were highlighted with triangles along the diagonal of the contact map. Key epigenetic regulators including CTCF, POL2RA, H3K4me3, H3K4me1, and H3K27ac were illustrated with the colors, black, magenta, and orange. The color, cyan represents H1 cells and the color purple represents GM12878 cells. Pink triangles denote transcription start sites (TSS). **D**. The heatmap illustrating the number of six types of NucIL, NucD-NucD, NucD-NucB, NucB-NucB, NucD-NucG, NucB-NucG, and NucG-NucG in GM12878 cells. **E**. The bar plot presenting the number of NucILs across different classifications: total NucILs, promoter-related NucILs or P-centric NucL (P1 NucIL), promoter-enhancer interaction type 1 related NucILs (P1+EI-T1 NucIL), and the associated genes for these categories in H1 and GM12878 cells (P1+EI-T1 genes). **F**. The Venn diagram showing the number of P1+EI-T1 genes that are H1-specific, common between H1 and GM12878, and GM12878-specific. **G**. The top biological pathways enriched with H1-specific and GM12878-specifc P1+EI-T1 genes. **H**. The nucleosome-based contact maps and integrated genomics viewer (IGV) tracks for a GM12878-specific gene, SPI1 genes in H1 and GM12878 cells, respectively. NucD regions were highlighted with triangles along the diagonal of contact map. The rectangles in the contact map showed the identified NucIL in each cell type. Key epigenetic regulators including CTCF, POL2RA, H3K4me3, H3K4me1, and H3K27ac were illustrated, with the colors, black, magenta, and orange. The color, cyan represents H1 cells and purple represents GM12878 cells. The arch line represents a P2P1 NucL between SPI1 and PSMC3.

Further comparison of NucILs between H1 and GM12878 cells revealed that although the total number of NucILs and the number of promoter-associated NucILs (P1 NucIL) were similar in the two cell types, both the number of EI-T1 within P1 NucILs (P1+EI-T1) and their associated genes were lower in GM12878 cells than in H1 cells (**Fig. 6E**,**F**). This was in line with an observation that there are fewer expressed genes in GM12878 than in H1 cells (**Extended Data Fig. 8I**). Intriguingly, H1-specific P1+EI-T1 NucIL-associated genes were enriched in development-related pathways (**Fig. 6G** – top) while GM12878-specific ones were enriched in lymphoblastoid-related pathways (**Fig. 6G** – bottom), underscoring Nuc3DMap’s capacity to identify cell-type specific interaction loci, exemplified by the SPI1 gene shown in **Fig. 6H**. In summary, we have demonstrated that Nuc3DMap is a robust algorithm and novel computational tool to identify nucleosome-based chromatin interactions efficiently and effectively in a cell-type-specific manner.

## Discussion

Here, we developed a novel algorithm and computational method, Nuc3DMap, to comprehensively integrate MNase-seq, Hi-C and Micro-C data. To the best of our knowledge, our Nuc3DMap is the first computational tool designed to delineate finer nucleosome-based 3D chromatin architecture at multiple levels. We rigorously tested and evaluated it on publicly available MNase-seq, Hi-C and Micro-C data in H1 and GM12878 cells, respectively, and identified nucleosome-based chromatin interactions in cell-type-specific manner, including H1-specific or GM12878-specific NucDs, NucBs and NucLs. These NucBs and NucLs can be further classified into distinct functional types by integrating hundreds of cell-type-specific combinatorial transcriptional and epigenetic regulators using hierarchical and autoencoder-GMM methods. Interestingly, functional NucBs were associated with distinct nucleosome organization and exhibited varying insulation capabilities, suggesting an epigenetic context-dependent insulation ability of topological boundaries. Intriguingly, we identified a type of functional Pol2-related NucLs that were involved in cell-type-specific promoter-enhancer communications, indicating a noncanonical regulatory role of Pol2 in mediating gene looping and cell type specificity.

Surprisingly, only a small portion of NucBs were associated with one or two more of five STFs, CTCF, RAD21, SMC3, YY1 and ZNF143, in either cell types (**Fig. 4B and Extended Data Fig. 8B**), suggesting that many finer nucleosome-based topological boundaries may be involved in distinct regulatory functions rather than act simply as structural insulation. As a matter of fact, the majority of NucBs in either cell type was associated with one of TransRs or EpiRs including 26 HIS, 36 TF, 3 CR, 12 HISase, R-Loop, 5mC, 5hmC and nasRNA, and with a distinct type of functional NucBs, implicating its specific regulatory and epigenetic roles. Interestingly, some of NucBs seem to have a role in regulating cell type specificity (**Fig. 6C**). Our observations are consistent with the results from other recent Micro-C studies such that fine-scale chromatin organization is linked with gene activity, transcriptional regulation, and gene silencing^9,70^.

Intriguingly, our quantitative modeling of the relationship between P-centric NucL intensity and gene expression identified four distinct classes associated with different clusters, suggesting that specific TranRs and EpiRs might regulate the gene expression within each of four classes, EI-T1, EI-T2, EI-T3, and EI-T4 (**Fig. 5I**). We speculate that the class EI-T1 might be associated with cell type specificity. Interestingly, our data are consistent with recent studies showing the expression-distance scaling relationship or long-range activation decays with the E-P genomic distance by using CARGO-VPR^71^. Taken together, our work systematically demonstrates the relationship between interaction intensity and transcriptional activity, unveiling an activation-like behavior of chromatin looping.

There are several notable strengths of Nuc3DMap. Firstly, Nuc3DMap constructs a nucleosome-based contact matrix which can preserve nucleosome-free regions and offer a precise mapping between high-order chromatin structures and linear genomic features. Unlike typical fixed-bin segmentation, Nuc3DMap incorporates nucleosome positioning information during constructing contact maps, allowing a more accurate delineation of finer chromatin architecture such as nucleosome-based topologically associating domains and nucleosome-based interaction loci. Secondly, Nuc3DMap implements an advanced computing technique, a sparse Knight-Ruiz algorithm, to ensure robust and efficient balancing of extremely sparse and large-scale contact matrices. This technical advancement mitigates a limitation of handling large and sparse chromatin interaction datasets, improving the accuracy of nucleosome-based contact inference. Thirdly, Nuc3DMap leverages a bottom-up renormalization grouping approach which allows investigation of chromatin architecture at multiple scales while preserving the ability to trace back to the fundamental DNA packaging unit, the nucleosome. This provides a more precise and targeted connection between high-order 3D chromatin structures and the linear epigenome. Lastly, Nuc3DMap provides a foundational framework, *i*.*e*., a nucleosome-based contact matrix, to the broader research community so that more computational tools based upon this matrix are developed for chromatin organization analysis. For instance, users can leverage Nuc3DMap to refine or expand the identification of nucleosome-based chromatin features, including but not limited to compartments, TADs, loops, stripes, and spandrels^72^. This flexibility positions Nuc3DMap as a powerful resource for advancing computational methods in dissecting nano-scale 3D genome organization, enabling new discoveries and fostering innovation in chromatin structure modeling.

Despite its strengths, Nuc3DMap has serval limitations. Its reliance on nucleosome-resolution contact matrices necessitates deep sequencing coverage, requiring at least 1 billion high-quality, non-duplicated pairs for both Micro-C and Hi-C to accurately capture nucleosome-based chromatin architecture, such as NucDom and NucIL. While this requirement may present a financial challenge, particularly for large-scale studies or clinical applications, our robust testing indicates that increased sequencing depth significantly enhances the accuracy of chromatin architecture reconstruction. In this regard, the high data requirement is not merely a limitation but an opportunity for providing richer, more precise insights into nucleosome-based chromatin organization. Additionally, while Nuc3DMap successfully integrates nucleosome occupancy and chromatin interaction data, it does not currently incorporate nucleosome orientation information, limiting its ability to resolve directional nucleosome positioning along chromatin fibers. This feature has recently been reported as essential for pioneer factor binding^73^. Future integration of datasets such as Hi-CO, which provides orientation-aware chromatin interactions, could further improve the resolution and functional interpretation of nucleosome interactions. Furthermore, structural variations, a key factor in genome organization, are not yet accounted for in the current implementation of Nuc3DMap, representing an area for future development. Finally, inter-chromosomal interactions are currently not implemented due to the computational challenges associated with handling excessively large matrices. Future efforts could be made to require a batch-processing approach to ensure proper normalization and maintain computational efficiency while accurately capturing inter-chromosomal interactions.

Overall, Nuc3DMap represents a conceptual advancement in chromatin conformation analysis, offering a refined view of genome organization from the fundamental nucleosome level to higher-order chromatin structures. By bridging nucleosome-level resolution with broader chromatin architecture, Nuc3DMap provides an unprecedented framework for understanding chromatin folding dynamics and regulatory complexity. By delivering a high-resolution, nucleosome-based perspective on genome organization, Nuc3DMap has the potential to drive new discoveries in chromatin biology and gene regulation across diverse cellular contexts, ultimately contributing to a more comprehensive understanding of genome function in health and disease.

## Methods

### The detailed algorithmic implementation of NucPrep and NucLoad modules

In Nuc3DMap, there are five modules, including: NucPrep, NucLoad, NucMerge, NucDom, and NucIL. A detailed algorithmic implementation of each of five modules was described in the following.

The NucPrep module: Pre-processing MNase-seq, Hi-C and Micro-C data into nucleosome locations and chromatin interaction pairs by using publicly available tools and making the files compatible formats (.bed and .pairs) for NucLoad module. For MNsas-seq data, NucHMM^19^ was used to derive the nucleosome locations. For Hi-C and Micro-C data, the Dovetail Genomics pipeline (https://micro-c.readthedocs.io/en/latest/fastq_to_bam.html) was adapted to identify chromatin interaction pairs. The pipeline was modified using pairtools^41^ parse2 by adding --add-columns pos5, pos3 and --report-orientation read parameters. These two modifications are essential since the nucleosome locations binarize the genome into variable-sizes bins, so that both genomic coordinates of the bins are necessary to precisely associate chromatin interaction pairs with the nucleosome-based bins (**Fig. 2E**).

The NucLoad module: Building Hi-C and Micro-C nucleosome-based contact matrix, respectively. It consists of the following seven steps (**Extended Data Fig. 1**):

*Step 1: Genome nucleosome-binarization*: The first and key step to build a nucleosome-based contact matrix is the binarization of the whole genome based on the nucleosomes and the genomic regions between nucleosomes. Here, we defined two types of bins in the nucleosome-based binarization: N-bin and I-bin, where N represents nucleosome, and I represents a genomic region between two index-adjacent nucleosomes. We applied the following strategies to obtain all Ns and Is: For Ns, we first acquired all potential nucleosome core locations from NucPrep, then filtered out ‘shoulder’ nucleosome cores which are the edge signal for nucleosomes^40^:

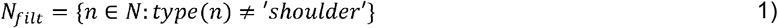

We further extended nucleosome cores to 150 bp for those nucleosome cores shorter than 150 bp and kept their original length for those nucleosome cores longer than 150 bp:

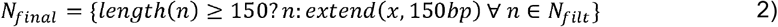

For Is, we first picked all regions between two index-adjacent nucleosomes as the potential intervals, then filtered out the regions shorter than half standard nucleosome size (75 bp):

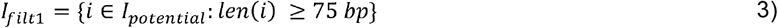

I *I* filt_1_ was further filtered out based on Eq 4 and 5:

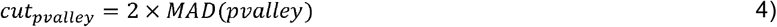

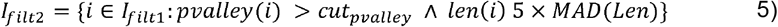

*where* MAD *is the median absolute deviation*. This filtration step was designed to exclude intervals that were relatively short and lack sharp MNase-seq peaks flanking around them.

Lastly, we utilized the following rules to construct nucleosome-based intervals from the filtered intervals: 1. regions with length shorter than two standard nucleosome size (300 bp) were defined as an Interval; 2. regions with length longer than 300 bp but less than 600 bp were equally divided into two Intervals; 3. regions with length longer than 600 bp less than 30,000 bp were divided by 300 bp but the last two bins was calculated by mean length; and 4. regions with length longer than 30,000 bp were equally divided into 100 bins, denoted as:

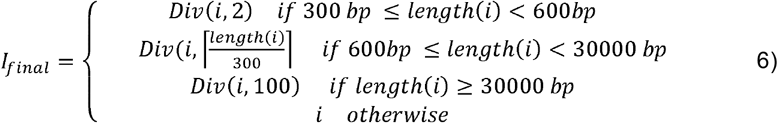

In the end, a contact matrix is the combination of filtered N-bins and processed I-bins. For the convenience of description, N and I were collectively denoted as B={*b*_*1*_, *b*_*2*_, … *b*_*i*_, … *b*_*R*_}, *i* ∈ {∀ *Nucleosome & Intervals*}, respectively, and the total number of bins as R.

*Step 2: Initialization of the contact matrix*: After nucleosome-based binarizing the whole genome, we then initialized a R-by-R nucleosome-based empty contact matrix, denote as *C*_null_.

*Step 3: Partition of interaction pairs into Loci list*: A chromatin interaction pair was partitioned into two lists, *L*_1_ and *L*_2_, where each list contained loci from one end of the chromatin interaction pairs. Each of the elements in *L*_1_ and *L*_2_ was in a one-to-one correspondence and indexed as *L*_1_ [*i*] and *L*_2_[*i*] indicating they were from the same interaction pair.

*Step 4: Application of NCLS Algorithm*: We applied the Nested Containment List (NCLS) algorithm implemented in PyRanges^74^ to efficiently identify overlaps between loci and nucleosome-base bins. For each pair of interaction loci (*L*_1_ [*i*] and *L*_2_[*i*]), the NCLS algorithm was able to find all bins that overlap with at least one base pair. This overlap was determined by the following Eq 7:

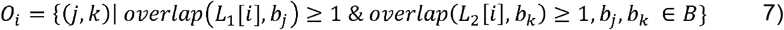

*where overlap function finds the bases overlapped between two genomic coordinates*.

*Step 5: Filtering ‘random’ overlaps*: After is determined, a filtering step is to remove ‘random’ overlaps using an overlap cutoff determined by MAD of overlap lengths. To enhance flexibility, users can also manually define this cutoff value.

*Step 6: Resolving multiple binning conflicts*: If a loci end overlaps multiple bins, the bin with the greatest overlap was identified. Optionally, users can choose to skip this resolution to allow an end to be assigned to multiple bins, facilitating multi-assignments.

*Step 7: Filling contact matrix*: For each identified pair (*j*,*k*) ∈ 0_*i*_, the corresponding matrix entry *C*_*j*,*k*_ was incremented by 1. This step generates accumulated contact matrix *C* from *C*_*null*_

### Merging normalized Micro-C and Hi-C nucleosome-based contact matrices

The NucMerge module consisted of four major steps: norm-anchor identification, margin adjustment, sparse matrix balancing and contact matrix merging.

*Step 1: Norm-anchor identification*: This initial step was involved with the identification of normalization anchors based on enzyme digestion sites that were consistent across different datasets. These anchors were critical for aligning interaction pairs from Hi-C and Micro-C data and served as reference points for subsequent normalization. The norm-anchors were identified with the following equation:

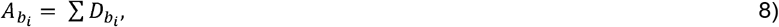

*where* 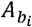 *is the anchor score for b*_*i*_ *and D is the presence of a digestion site at b*_*i*_. *b*_*i*_ *with A*_*i*_ ≥ 1 *are identified as norm-anchor for further adjustment*.

*Step 2: Margin adjustment*: Margin adjustment was performed to align the margins of the Hi-C and Micro-C matrices, ensuring that the sums of rows and columns across the datasets were comparable. This adjustment was essential for correcting global differences in library size or sequencing depth. For the convenience of description, we denoted margin value of Micro-C contact matrix for each *b*_*i*_ as 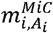 and margin value of Hi-C contact matrix for each *b*_*i*_ as 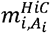. Detailly, we first constructed a list of tuple 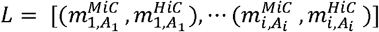 containing paired margin value from Micro-C and Hi-C nucleosome-based contact matrix. We then separated *L* into different groups {*L*_*1*_, *L*_*2*_, …, *L*_*Z*_} based on anchor score, where Z is max ({*A*_1_, …, *A*_*N*_}). For each *L*_z_, we filtered out outliers with a modified Z-score larger than 3.5^75^, where the modified Z-score was calculated by

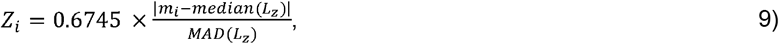

Then, 1-degree polynomial fit was applied on filtered *L*_z_ and polynomial coefficient 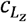 was calculated for each filtered *L*_z_. Based on 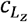, we screened out *L*_z_ with large discrepancy 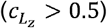 between Micro-C and Hi-C data for further normalization. We denoted *L*^*filt*^ as the new list contained all 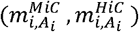 that met all above criterion. For the convenience of calculation, we converted *L*^*filt*^ into a two-column matrix, denoted as 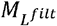, and performed log transformation on 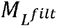:

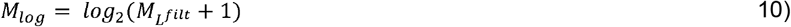

We then normalized *M*_*log*_ with the following Eq 11:

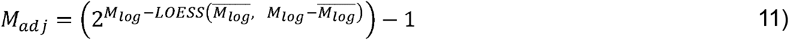

*where, LOESS represents locally weighted polynomial regression and* 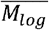 *represents row mean of M*_l*og*_.

We finally performed a linear interpolation for the rest of *b*_*i*_ based on the *M*_*adj*_ and acquired adjusted margin value of eachi *b*_*i*_, denoted as *NM*_*i*_.

*Step 3: Sparse matrix balancing*: After margin adjustment, we needed to balance the original matrix with the restricted margins. As our matrix was extremely sparse and large, we performed an adapted KR algorithm, sparse-KR (sKR), to achieve the restricted sparse matrix balancing. sKR consisted of two major steps: matrix pre-processing and restricted matrix balancing. The pre-processing step speeds up the balancing process and makes sure the matrix has total support (row permutations exist which allow for every nonzero element in the matrix to lie on a main diagonal that contains no zeros), which guarantees the existence of a scaling. Restricted matrix balancing scaled the original Micro-C and Hi-C contact matrix based on the normalized margin value. More specifically, zeros rows, *i*.*e*., all elements in the row are zero, were first removed from the contact matrix *C* to give *C* ′, where *C* ′= *C* [*i*,:] for all i such that ∑*j C*[*i*,*j*] ≠ 0, and *C* [*i*,:] *represents the i-th row of C*. The matrix *C* ′ was then reordered into a block diagonal structure by identifying connected components and reordering accordingly :

Let *C*_*1*_, *C*_*2*_, …, *C*_*k*_ be the connected components identified in the graph, the matrix *C* ′ is transformed into *T* such that

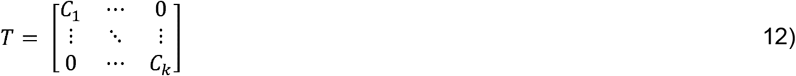

which allows for faster scaling. For each block, we added *ϵI* to ensure the total support (If the block could be balanced without this adjustment, then the loss of fidelity is no bigger than □), the adju sted block 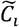 is given by

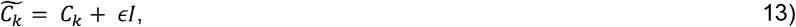

*where I is the identity matrix of the same dimension as C*_*i*_, and *ϵ is a small positive scalar (practically given by numpy*.*finfo*.*eps*).

For each 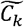, we refined the KR algorithm^76^ to balance the matrix while targeting a specific margin value. The goal is to find positive vectors *γ* and *c* such that

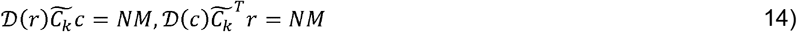

Since our contact matrix is symmetric, then the above equation can be simplified by letting *r* = *c*. To achieve balancing, we need a weight vector *w*_*_ that satisfies

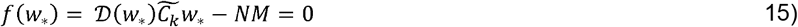

We then iteratively adjusted *w*_*_ to minimize the difference between the row/column sums of *𝒟*( *w*_*_) *Aw*_*_ and *NM* with

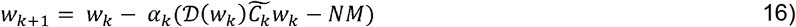

*where* 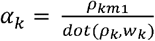 *is the step size for the iteration calculated to ensure convergence, ρ*_*km*1_*is a measure of the residual from the previous iteration*, and *dot* (*ρ*_*k*,_ *w*_*k*_) *is the dot product of the update direction and the matrix-vector product involving the current estimate w*_*k*_. This formulation adaptively scales *α*_*k*_ based on how effectively the previous update reduced the residual.

After *w*_*_ is measured for all blocks, we combined all *w*_*_ into a single weight vector *w*. We then traced back the index in *T* and re-ordered the *w* to form *w′*. For zeros rows, we inserted zero weight into *w′*. This process allowed us to generate a scaling vector for each *b*_*i*_. Finally, given a scaling *W*, the Micro-C and Hi-C contact matrix was balanced by the simple multiplication:

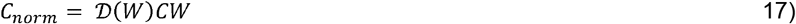

*Step 4: Contact matrix merging*: After Micro-C and Hi-C nucleosome-based contact matrices were balanced, we merged two contact matrices by using Micro-C contact matrix as the base matrix to incorporate Hi-C unique pixels. Briefly, we first filtered pixels in Hi-C contact matrix associating with zero anchor score to reduce the noise and keep more reliable interactions:

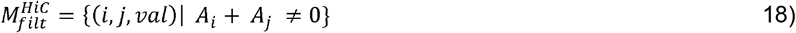

We then determined the unique Hi-C interactions by searching the pixels with non-zero values in 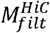 but with zero values in 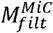 :

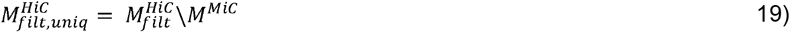

The merge contact matrix is the summation of the Micro-C contact matrix and Hi-C contact matrix.

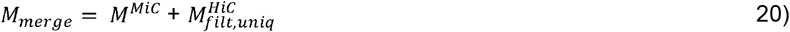

sKR algorithm was finally applied again with (all elements are one) as the margin value to adjust the contact frequencies. This step was to correct systematic biases and variations in interaction frequencies across the genome as describe in Rao et. Al^6^.

### Detecting NucD, NucB, NucG on the merged contact matrix

The NucDom module was designed to identify nucleosome-based topological domain, boundary, and gap, termed as NucD, NucB and NucG. It adapted the main workflow of TopDom^29^ with modifications that accommodate variable-sized and discontinuous binarization on forming our nucleosome-based contact matrix. NucDom has three steps: firstly, a signal vector (V) was calculated by averaging a sliding diamond-shaped window across the *b*_*i*_

Let *s* denote the size of the window and *m*_*i*,*j*_ represents the contact frequency of i-th row and j-th column in *M*_*merge*_, each element *V*_*i*_ in V can be measured with:

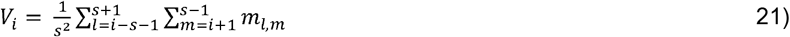

After V was calculated, turning points (tp) and local extremes were detected by curving fitting algorithm: 1. Initialize the feature value array *F*_*ν*_ with *F*_*ν*_ [0] =0 and the error value array *E*_*ν*_; 2. For each of two elements *V*_*i*_ and *V*_*j*_, *j* >*i*,

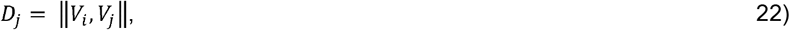

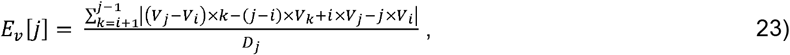

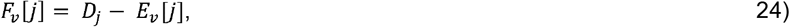

The tp were identified where *F*_*ν*_[*j*] decrease from *i* to *i* +1; local extremes were further classified from tp, *i*.*e*., Local maximum 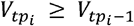 and 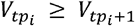 and Local maximum 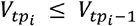 and 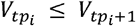 Then, a Mann-Whitney test was used to assess whether there is a significant difference between within the up- or downstream windows and interactions between the up- and downstream windows for each *b*_*i*_ as described previously^21^. False positive boundaries were then filtered, and scores were assigned to each identified domains/boundaries and gaps based on the scaling p value.

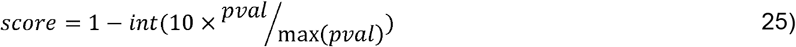

After identifying NucD, NucB, and NucG (collectively referred to as features), each of these features was mapped back to specific nucleosome indices, in which it linked high-level chromosomal architecture directly to the underlying nucleosome structure, which is also vital for visualizing and detecting nucleosome-based interaction loci. Lastly, the genomic coordinates in the output file were consecutive without overlaps and gaps between adjacent features.

### Identifying NucIL from the merged contact matrix

The NucIL module was to identify significant nucleosome-based chromatin interaction loci (NucIL). The process was structured into two main steps in the following:

*Step 1: Renormalization group matrix construction*: As the sparsity of the original merged nucleosome-based contact matrix is very high, we applied renormalization group approach to construct a renormalization group (RG) matrix, denoted as *C*^*RG*^ to reduce the sparsity and increase signal-to-noise ratio as described: Firstly, we scaled elements in the merged contact matrix based on the decay spline between distance and contact frequency. Detailly, let *c*_*i*,*j*_, denoted the pixel value located at i-th row and j-th column, which represent the contact frequency between *b*_*i*_ and *b*_*j*_. The distance of the pixel was then calculated as

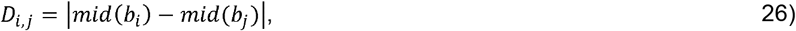

*where mid (b) is the midpoint of bin b*.

Then, all *c*_*i*,*j*_ were sorted by *D*_*i*,*j*_, denoted the sorted list as *SC*. We then divided the *SC* into 200 bins, with each bin containing the same number of *c*_*i*,*j*_:

Let *SC* ={*sc*_*1*_, *sc*_*2*_, … *sc*_*k*_}, *k* ≤ 200, and we then calculated average contact frequency for each *sc*_*k*_;

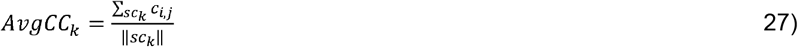

*where* ‖ *sc*_*k*_ ‖ *represents the number of c*_*i*,*j*_*in sc*_*k*_. For each *sc*_*k*_, we also measured the range of *D*_*i*,*j*_, within it, denoted as [*d*_*max*,*k*,_ *d*_*max*,*k*_]. Then, *AvgCC* was fitted by UnivariateSpline in Scipy^77^ with *splineErr*=*min*(*AvgCC*)^2^, and *DistCoef*_*k*_ was calculated by fitting the spline. Finally, we could scale *c*_*i*,*j*_ into 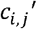 with:

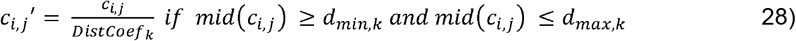

Given a set of genomic bins *B =*{*b, b*_*2*_, …, *b*_*N*_}, based on the identified NucD/NucB/NucG, we categorized these bins into distinct regions, represented as *R =*{*r*_*1*_, *r*_*2*_, …, *r*_*U*_}, where *U* is the total number of unique regions defined by NucD, NucB, and NucG. Each region *r*_*u*_ is a subset of *B* containing bins that are grouped together based on their NucD annotation.

The renormalized contact matrix *C* ^*RG*^ was then constructed by aggregating contacts from the original matrix *C* based *on R with* block size correction, Specifically, the element 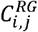 in the renormalized matrix was computed as follows:

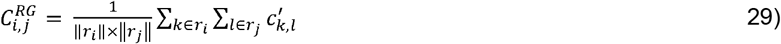

*where r*_*i*_ and *r*_*j*_ *are sets of indices corresponding to the bins grouped under the regions i and j* respectively. And ‖*r‖ is the size of r*.

By applying this RG approach, the dimension of the contact matrix reduced ~100x, e.g, RG matrix for chr1 in H1 is 91,461 x 91,461, while original contact matrix is 1,127,835 x 1,127,835. *Step 2: NucIL detection*: We applied a dual-step to detect NucIL, including using the SIFT algorithm described in MUSTACHE^32^ to capture significant interaction loci and using quantile detection (QD) to identify high-intensity interaction loci failed to be detected in the SIFT step. The SIFT detection comprised three steps: matrix pre-processing, applying the Difference-of-Gaussian (DoG), and feature localization with refinement. Initially, we divided RG matrix of the chromosome into smaller chunks based on the genomic distance, which helped to restrict the distance range of interaction loci, 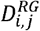, and to manage the computational load. We then removed low-value pixels in the matrix and applied a triangular upper matrix form to avoid potential duplication. For DoG step, let *ch*_*g*_ be g-th chunk of the triangular upper RG matrix. The gaussian smoothing performed on the *ch*_*g*_ was defined:

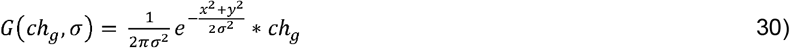

*where* * *denote the convolution operation, is the standard deviation of the Gaussian kernel for controlling the extent of smoothing which can be customized*. The DoG was defined by calculating difference between two adjacent *G*(*ch*_*g*_, *σ*):

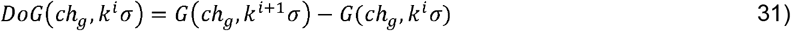

*where k is constant that scales the standard deviation for adjusting the difference in smoothing levels between the two applications of the Gaussian filter*.

To detect NucIL in each genomic chunk, *ch*_*g*_ underwent a series of Gaussian smoothing at progressively increasing scales. These scales were geometrically spaced by a constant factor *k*, {*σ*,*kσ, k*^*2*^*σ*,… *k*^*s*^*σ*}, where s is the customized input parameter for controlling the level of scales. This produced a sequence of smoothed contact maps for each chunk, each corresponding to a different level of detail or scale. The sequence, often referred to as octaves, consists of each map being a blurred version of the original, with the degree of blurring increasing. The DoGs were then calculated by subtracting each pair of adjacent *G*(*ch*_*g*_, *σ*). This subtraction amplified regions where interactions become pronounced or diminished significantly as the level of smoothing changes, thus pinpointing potential interaction loci. The potential local maxima were identified on DoG with:

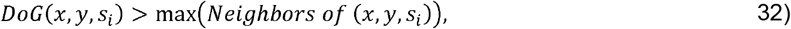

*where s*_*i*_ *represents a specific scale*, and *‘Neighbors’ includes the eight surrounding pixels at scale s*_*i*_ *plus the corresponding neighbors at scales s*_*i*+1_ *and s*_*i*-1_. Each local maximum underwent a validation against a significance threshold using p-values. These p-values were derived from:

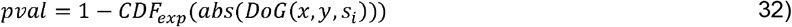

*where CDF*_exp_ *is the cumulative distribution function of the exponential distribution fit to the absolute values of the DoG responses*. Features that passed the initial checks were further refined by connectivity and sparsity as describe in MUSTACHE.

We next extended the framework to incorporate QD. This subsequent step was crucial for capturing significant interactions that may not have been sufficiently emphasized by SIFT alone (due to connectivity or sparsity, etc), thereby ensuring a comprehensive detection of interaction loci from the RG matrix. Similar to the SIFT method, we first filtered pixels by genomic distance to focus on interactions within specific bounds (default 2 Mb). Then, the intensities of contacts were then subjected to a quantile cutoff by computing the 0.999 quantile:

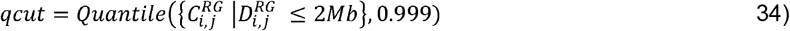

Only interaction loci with intensity values above this threshold were considered as significant candidate loci which were further grouped. This grouping used a window size to group all interaction loci within the window together and these grouped loci were the highest intensity of interaction. Finally, the NucIL detected from both SIFT and QD steps were combined upon removing the duplicates and grouping the adjacent pixels.

### Definition of genomic location

We defined five genomic locations as followed: A desert region was the region extending beyond 250 kb upstream or downstream of a transcription start site (TSS) of any annotated genes. A distal region was defined as the intermediate area located between 10 kb and 250 kb from a TSS, both upstream and downstream. A proximal region was the region extending from 1 kb to 10 kb, both upstream and downstream from a TSS. A promoter was the regions spanning 1 kb upstream to 1 kb downstream of a TSS. A gene-body included the entire gene region from start to end.

### Classification of Intervals

Intervals were systematically classified into four categories based on their genomic location length and MNase-seq signal intensity (determined by pvalley feature from iNPS): nucleosome-free regions (NFRs), nucleosome low-intensity regions (NLIRs), nucleosome-depleted gaps (NDGs), and desert regions (DRs). NFRs were the intervals with very low MNase signals and predominantly found in promoters with no size restriction. Additionally, intervals smaller than 600 bp located in distal or proximal regions or within gene bodies were also categorized as NFRs. NLIRs were characterized by low to middle level of MNas signals, i.e., flattened flanked MNase-seq signal peaks with middle-level pvalley value. NLIRs were located in any genomic regions except deserts. NDGs were located within the distal, proximal, or gene body regions and must exceed a size threshold of 600 bp with very low MNase signals, *i*.*e*., sharp flanked MNase peak signal with large pvalley value. DRs were characterized either by their location in the desert regions of protein-coding genes or by their size which exceeds 5 kb.

### Comparison of NucLoad, Bedtools and iNucs

To evaluate the performance of NucLoad, we conducted a comparative analysis against two established tools: Bedtools^78^ and iNucs^20^. Our experimental setup involved constructing datasets comprising various pair counts, ranging from 1 million to 9 million pairs. In these datasets, 90% of the pairs were selected based on both ends overlapping with a nucleosome by at least 1 bp, while the remaining 10% were selected by ensuring both ends overlap with predefined intervals, thereby introducing noise into the data. For a fair comparison across all tools in terms of input reading and output writing, we utilized a custom script that maintained consistency in reading input pairs and writing outputs in COO format (coordinate list format). Each of three tools were operated on an identical nucleosome location file generated from NucPrep. The specific procedure for each of three tools was the following: for NucLoad: execution followed the previously described method; for Bedtools: a custom script utilized the intersectBed function to detect overlapping pairs; for iNucs: the procedure followed the manual from its GitHub repository using default settings. Two performance metrics were evaluated. Speed was measured using the Python time package and reported as the time taken (in seconds) by each package to produce outputs from given inputs. Accuracy was defined by the count of reads correctly assigned to nucleosomes. For NucLoad, both best match mode (default) and multiple mapping mode were tested, as Bedtools script aligns more closely with the multiple mapping mode, providing a comprehensive comparison across different operational settings.

### Parameter optimization for NucDom and NucIL

We performed a parameter optimization for both NucDom and NucIL. NucDom’s functionality depends on a single parameter, Winsize. We tested NucDom across 18 different Winsize ranging from 3 to 20. For each Winsize, we measured the number of CTCF peaks located at the NucB. Further, we examined the contact frequencies within NucD (intra-NucD) *v*.*s* between NucD (inter-NucD, *i*.*e*., intra-NucB or NucG). The contact frequency difference between intra-NucD and inter-NucD was calculated as follows:

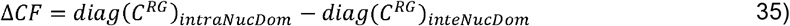

*where*, diag *is a function to acquire the diagonal of the matrix and C*^RG^ *is renormalization grouping matrix as mentioned above*. To comprehensively evaluate the performance of each Winsize configuration, we devised a simple scoring system. This score was computed by summing the ranks, where the ranks were assigned in descending order based on the count of CTCF peaks at NucBs and the median value of the intra-inter contact frequency discrepancy. The Winsize that yielded the highest cumulative score was selected as the optimal parameter.

Since NucIL has more parameters, we adopted the number of scales and octaves from MUSTACHE by setting them to 10 and 2, respectively. We particularly evaluated three main NucIL tunable parameters: the sparsity threshold (st), the p-value threshold (pt), and the initial sigma (sigma0). We tested a total of 64 combinations (4×4×4) of these parameters, st values of [0.25, 0.5, 0.75, 1.0], pt values of [0.01, 0.05, 0.1, 0.2], and sigma0 values of [0.25, 0.5, 0.75, 1.0]. Our evaluation metrics included: 1) the number of detected NucILs; 2) the length of NucIL (distance between two loci); 3) the intensity of NucIL and 4) aggregated peak analysis (APA) score. When st and sigma0 were set to 1.0, the number of NucIL and its length reduced significantly. A sigma0 of 0.75 resulted in the lowest median intensity of NucILs. higher pt values consistently produced more NucILs with lower median intensity. In the absence of a gold-standard criterion, we empirically set the default pt at 0.1, sigma0 at 0.5, and internally using a combined result from st 0.25 and 0.75.

### Robust Test

To accommodate a nucleosome-level resolution of the Nuc3DMap contact matrix, a higher sequencing depth is required compared to a traditional kilobase-resolution matrix. We performed a robust test to investigate the impact of the sequence depth to the NucDom and NucIL and to determine the minimum sequence depth for Nuc3DMap. We constructed a series of Micro-C datasets at varying proportions of the original dataset size, 25% (m0.25), 50% (m0.5), 75% (m0.75), and 100% (m1). A similar process was done for a series of Hi-C datasets. This resulted in 16 unique combinations (4 Micro-C × 4 Hi-C) with different sequencing depths. For each combination, we applied NucLoad, NucMerge, NucDom and NucIL and counted the number of NucDs, NucBs, and NucGs across all combinations. Additionally, the preservation of domain coordinates (identical start and end positions) was assessed. We further explored the genomic correlation between datasets by transforming the whole genome into a vector representing: 1 (NucD), 0 (NucB), and −1 (NucG). For instance, the vector [1, 1, 1, 1, 1, 0, 0, 0, −1, −1, −1] indicates that the first five base pairs were identified as NucD, followed by three base pairs as NucB, and the last three as NucG. We then calculated the Pearson correlation coefficient between any two conditions.

### Classification of NucB

We curated a wealth of TR data from multiple sources, including histone marks, TFs, 5mC, 5hmC, R-loops, and nascent RNA (**Extended Data Table 1**)^69,79–81^, Peak files for each dataset were primarily obtained directly from these sources. In instances where peak files were unavailable, we generated peaks using MACS3 bdgpeakcall^82^ against the hg38 reference genome with parameters optimized via the ‘--cutoff-analysis’ option. We constructed a NucB-Peak matrix, *M*^*NucBP*^, where each row corresponds to a NucB and each column to a TR data. Overlapping between peaks and NucB extended with ±200 bp was identified using the Bedtools intersect function^12^, with the following criterion:

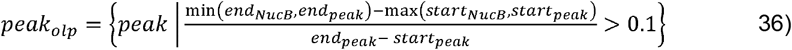

*where* olp *denotes overlap*. Each matrix element was then filled with the signalValue columns from the corresponding peak file. We further performed column-wised normalization of *M*^*NucBP*^ with capped Box-Cox transformation:

Let 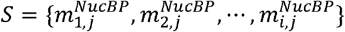 represents a column in *M*^*NucBP*^, we first capped S with 90^th^ quantile 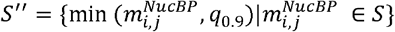; then all values were transformed

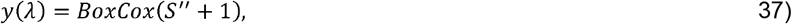

*where y*(*λ*) *is computed as*:

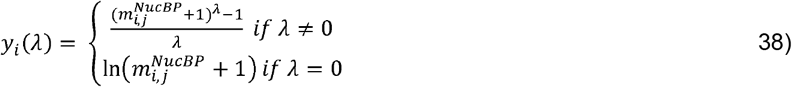

*where λ is determined internally by stat*.*boxcox function*. 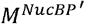 *is denoted as the transformed matrix*.

We then constructed a summarized matrix, denoted as 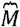, from 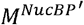 by incorporating 13 columns based on prior knowledge: A-HIS, R-HIS, P-HIS, E-HIS, Pol2, TFSS/TFNS, HISase, CR, STF, RLoop, 5mC, 5hmC and nasRNA. Specifically, the A-HIS and R-HIS columns contained the maximum values across their respective categories in 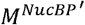 the P-HIS column was derived from the combination of A-HIS and R-HIS:

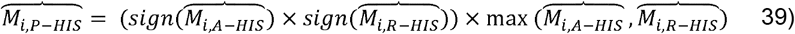

The E-HIS column contained the maximum value across all elongation histone modifications in 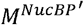; Pol2 column contained the maximum value among POL2RA and POL2RAphosS5 in 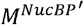; TFSS/TFNS columns contained the maximum value across all TFs with/without sequence-specific motif (except structural-related TFs) in 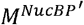; HISase contained the maximum value across all histone acetylation, deacetylation, or methylation complexes in 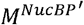; CR contained the maximum value across all ATP-dependent chromatin complexes in 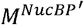; STF contained the maximum value across all five structural-related TFs, CTCF, RAD21, SMC3, YY1, ZNF143 in 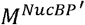; And RLoop, 5mC, 5hmC, as well as nasRNA contained the value of RLoop, 5mC, 5hmC, as well as nasRNA in 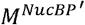, respectively. Upon constructing 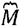, we further defined 12 combination types based on correlations observed between columns in 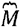 (**Fig. 4C**) and the frequency of NucB associated with each type (**Fig. 4D**). NucBs which were not classified within any of the 25 specified types are defined as “other” and have been excluded from further study.

### Nucleosome organization features for different types of NucB

We characterized nucleosome organization features, including nucleosome phasing, spacing, and positioning^19^, for different types of NucB. We first extracted nucleosomes and raw MNase-seq signals spanning a 2000 bp window centered on various NucB types using PyRanges and pyBigWig^83^, denoted as *W*_*NucB*_. For nucleosome positioning, we applied the equation as described^1^ for each nucleosome in *W*_*NucB*_

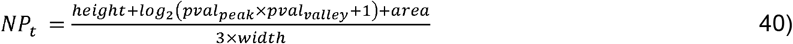

*where height, width, area, pval*_peak_ *and pval*_*valley*_ *are all from NucPrep*. We used the mean of all nucleosome positioning values for NucB types as the positioning score. For nucleosome spacing, we measured the distance between two adjacent nucleosomes in *W*_*NucB*_ (distances large than 500 bp was filtered out), and used the median of all nucleosomes spacing values for NucB types as the spacing score. For nucleosome phasing, following signal extraction, we aggregated the signals for each NucB type to obtain their cumulative profiles. The aggregated signal for each type was then normalized by dividing it by the number of instances within that type, followed by subtraction of the mean signal value across the region. This normalization process ensures that the signal reflects relative enrichment rather than absolute read counts, facilitating comparison across types. Additionally, we applied a Savitzky-Golay filter within Scipy^84^ to enhance the clarity of the signal and minimize noise. Lastly, we applied welch method described previous^81^ to calculate the phasing score from each NucB type signal. Specifically, we calculated the phasing score on left [−1000, 0] and right [0,1000] separately. We used the averaged phasing score from right and left as the final phasing score.

### Definition of nucleosome-based chromatin loop

We classified NucIL into gene-centric NucL based on the distinct genomic regions of its two loci. We defined three genomic regions, G (Gene-body), P (Promoter-Proximal), and D (Distal) as following: G: Spanning from 1000 bp downstream of the TSS to the transcription termination site (TTS). P: Defined as ±1000 bp from the transcription start site (TSS), extended to −5000 to 5000 bp. D: Encompassing ±250,000 bp from the gene boundaries but excluding P regions. Based on these definitions, NucL was systematically classified in the following: P2P1: both loci are located within the P regions of two different genes; G1G1: both loci are located within the G regions of the same gene; D1D1: both loci fall within the D regions of the same gene; D1P1: one locus is located in a P region and the other in a D region of the same gene; D1G1: one locus is within a G region and the other is in a D region; P1G1: one locus is positioned in a P region and the other in a G region of the same gene.

### Clustering of NucIL based on the transcriptional regulators and epigenetic regulators

To perform an unsupervised clustering of NucIL types, we curated a comprehensive dataset comprising seven TRs and seven histone marks from the ENCODE, including CTCF, RAD21, SMC3, YY1, ZNF143, POL2RA, TAF1, H3K4me3, H3K4me1, H3K27ac, H3K36me3, H3K79me2, H3K27me3, and H3K9me3. There were three steps in the clustering:

*Step 1: Data Extraction and Transformations*. For each instance of NucIL (*NucIL*_*i*_), corresponding to two loci denoted as *L*_*i*,1_ and *L*_*i*,2_, we extracted signal features from bigwig files corresponding to these specific genomic coordinates and computed both maximum and mean values for each TR and epigenetic mark at each locus. This computation yielded four 14-dimensional vectors per *NucIL*_*i*_:

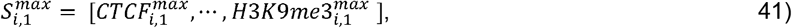

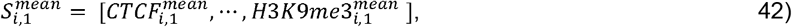

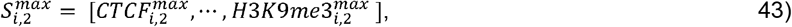

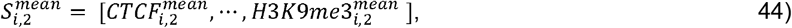

Upon the feature extraction, all vectors were normalized using the StandardScaler function from the sklearn library^22^ to facilitate a robust statistical analysis and ensure comparability across different samples and conditions. We formed a 28-dimension vector, *S*_*i*_, for each *NucIL*_*i*_ by averaging and combining four locus vectors:

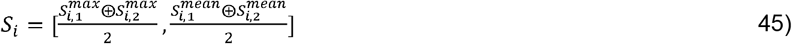

*where* ⊕ *represents element-wise addition*. We then constructed matrix ***X*** by aggregating all *S*_*i*_.*Step 2: Dimensionality Reduction and Clustering*. To determine the most effective method for dimensionality reduction, we initially established the optimal number of latent dimensions. This involved employing principal component analysis (PCA) to extract the principal components (PCs) that cumulatively explained up to 90% of the variance in the data. Specifically, PCA was configured to project the data ***X*** onto a lower-dimensional space represented by the eigenvector matrix, where each component’s significance was measured by its explained variance ratio. This method facilitated the selection of the top components necessary to capture the majority of the data variability. We then applied both PCA and an autoencoder (AE) for dimension reduction. PCA was performed to obtain latent space ***z*** from *X* by keeping first ten components. An autoencoder (AE) comprised an encoding function *f* that compressed the data *X* into a latent space ***z*** through a series of transformations:

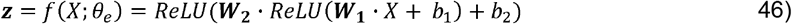

*where* ***W***_1_ and *b*_1_ are the weight and biases of the first linear layer mapping ℝ^28^ to ℝ^15^, and ***W***_2_ and *b*_2_ are the weight and biases of the first linear layer mapping ℝ^15^ to ℝ^10^, the latent space. The *decoder g* reconstructs the input data from the latent representation ***z***:

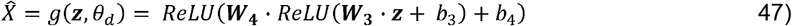

*where* ***W***_3_ and *b*_3_ expand ℝ^15^ to ℝ^10^, and ***W***_4_ and 4 reconstruct the original ℝ^28^ from ℝ^15^. The AE was trained to minimize the mean squared error (MSE) between the original and reconstructed data, quantifying the loss *ℒ* as:

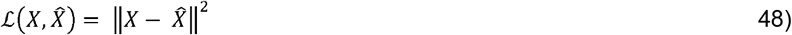

Optimization was performed using the Adam optimizer in order to reduce reconstruction error across the dataset:

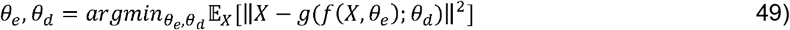

*Step 3: Optimal Clustering Selection*. After the dimensionality reduction, we implemented four clustering methods, PCA-KMeans, PCA-GMM, AE-KMeans, and AE-GMM, to select an optimal clustering. We evaluated each of these four methods on the number of clusters ranging from 10 to 20. We assessed PCA-KMeans and AE-KMeans using the within-cluster sum of squares (WCSS) and PCA-GMM and AE-GMM using the Bayesian Information Criterion (BIC). Both KMeans and GMM was implemented through sklearn. Additional two clustering evaluation algorithms, Calinski-Harabasz (CH) index and Davies-Bouldin (DB) index, were performed to further determine the number of clusters. Based on these metrics, we finally selected 14 clusters as the optimal number due to the lowest DB index score, which indicated the most distinct and well-separated clustering outcome.

### Nucleosome organization similarity

To calculate the similarity of the nucleosome organization of the cluster of NucIL, we first extracted and processed the MNase-seq signal from both loci and then measured Fréchet distance, denoted as frdist, as described in our previous study^19^. We finally converted frdist into nucleosome organization similarity between range [0,1] with the equation:

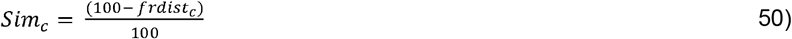

### Intensity-Expression curve

To investigate the relationship between P-centric NucL intensity and gene expression of each NucIL cluster, we initially scaled TPM obtained from ENCODE by adapting zFPKM method^86^: Let *E* be the input vector of gene expression values. Elements with zero values are excluded to prevent undefined logarithmic transformations, resulting in a modified vector *E* ′. The *E* ′ is then transformed using a base-2 logarithm: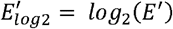 The final transformed was *E*_*z*_ was calculated:

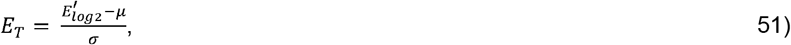

*where standard deviation* 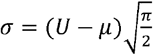 *is estimated from upper mean* (U) *and μ, incorporating an adjustment for half the width of the distribution*; *U is calculated as the mean of 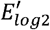 values greater than μ*; *and μ is the mode of distribution and identified as the point of maximum density in*

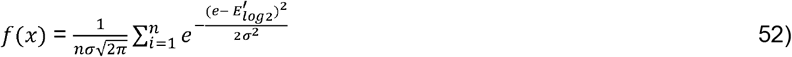

which is calculated over a specified range of 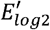. The *E*_*T*_ was segmented into 11 distinct classes using 10 quantile cutoffs, denoted as *q*_0.0_, *q*_0.1_, … *q*_1.0_ Each gene’s TPM value was then assigned a discrete label corresponding to its position within these quantiles. The labels were designated as integers ranging from −5 to 5 and denoted as *E*_*L*_. We then calculated the aggregated P-centric NucL intensity of each gene by summing the corresponded P1-related (see ‘***Definition of nucleosome-based chromatin loop’*** section) NucLs. We denoted *I* as the intensity vector for all gene. The *I* was further transformed to *I*_*T*_ by applying the Box-Cox transformation as described in ‘***The detailed algorithmic implementation of NucPrep and NucLoad modules***’ section. The *I*_*T*_ was also segmented into 11 distinct classes using 10 quantile cutoffs, denoted as *q*_0.0_, *q*_0.1_, … *q*_1.0_. Each gene’s intensity value was then assigned a discrete label corresponding to its position within these quantiles. The labels were also designated as integers ranging from −5 to 5 and denoted as *I*_*L*_.

For each cluster, we isolated the subset of data corresponding to that specific cluster from our previously processed data. A contingency table, 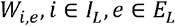, was then created using tallying the frequency of occurrences for each combination of *I*_*L*_ and *E*_*L*_. The intensity-expression curve *A* was derived from this table by following step: we assigned weight to each expression class based on its distance from the median class to emphasize the influence of classes that are farther from the median. These weights, *w*_*i*_ were assigned as

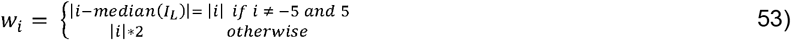

The intensity-expression curve for each intensity class was calculated as a weighted average of the transcription levels:

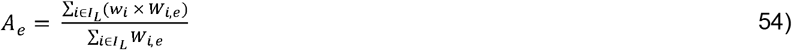

Lastly, we modeled the non-linear relationship between NucL intensity and gene expression level using a sigmoid function based on the observations of intensity-expression curves from different clusters of P-centric NucL. The function is defined as

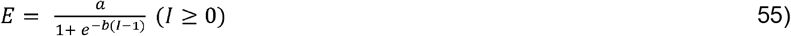

*where E* represents the expression level, *I* is the NucL intensity, *a* is a scaling parameter that adjusts the plateau value of *E*, and *b* is the steepness of the curve, controlling how quickly *E* increases/decrease with respect to.

### Comparison between H1 and GM12878 cells

We generated Micro-C data by our own and curated MNase-seq, Hi-C data, and various ChIP-seq datasets for GM12878 cells from public database^14^. Initially, we applied Nuc3DMap to identify NucD, NucB, NucG and NucIL in GM12878 cells. For the NucB, we classified it using the epigenomic data (**Extended Data Table 1**) to 19 NucB subtypes, including 11 single types, A-HIS, R-HIS, P-HIS, E-HIS, Pol2, TFSS/TFNS, HISase, CR, STF, 5mC, nascent RNA and eight combinatorial types, Active~STF, Poised~STF, Active~STF~nasRNA, Poised~STF~nasRNA, Active~Pol2~STF, Poised~Pol2~STF, Active~Pol2~STF~nasRNA, Poised~Pol2~STF~nasRNA. We conducted a comparative analysis of the NucB between H1 and GM12878 cells and defined Common (at least 1bp overlap between H1 and GM12878), H1-specific (exclusive to H1), and GM12878-specific (exclusive to GM12878) NucB. We then analyzed the frequency of the 19 NucB types. Subsequently, we employed the AE+GMM method to cluster NucILs in GM12878 cells, resulting in 16 distinct clusters. These were sequentially modeled and re-classified into four types: EI-T1, EI-T2, EI-T3, and EI-T4. We evaluated the total count of NucIL, the number associated with P regions (P1 NucL), those clustered as EI-T1 type associated with P regions (P1+EI-T1 NucL), and the corresponding genes in both H1 and GM12878 (P1+EI-T1 genes). We further identified H1-specific, GM12878-specific, and common P1+EI-T1 genes, and examined their GO pathways using DAVID, as previously described^87^.

### Visualization

We used a customized script to visualize the contact matrix based on matplotlib packages^88^ to enhance the plotting speed and adapted to variable-size bin. And other tracks (ChIP-seq, MNase-seq, R-Loop, 5mC, 5hmC and nascent RNA) were visualized by IGV^89^.

### Cell Culture

Human GM12878 B lymphocyte cells were obtained from the Coriell Institute for Medical Research. Cells were maintained in Roswell Park Memorial Institute (RPMI) 1640 Medium supplemented with 2 mM L-glutamine and 15% fetal bovine serum, cultured at 37°C in a humidified atmosphere containing 5% CO_2_. Cells were split every three days to ensure continuous growth and maintenance.

### Micro-C profiling

Micro-C profiling was conducted as previously described^9,13^. Briefly, 1 × 10 □ cells were harvested and subjected to cross-linking with DSG and formaldehyde at room temperature. Following cross-linking, chromatin was digested with MNase Enzyme Mix (Cat # PN DG-NUC-001, Dovetail Genomics, Scotts Valley, CA, USA) to achieve appropriate fragment sizes. Cells were subsequently lysed with SDS, and the lysate was mixed with Chromatin Capture Beads (Cat # PN DG-REF-001, Dovetail Genomics) for efficient chromatin binding. The mixture was incubated at room temperature for 10 minutes to facilitate chromatin attachment to the beads. End Polishing Master Mix (Cat # PN DG-NUC-001, Dovetail Genomics) was added to the chromatin-bound lysate. Incubation was performed at 22°C for 30 minutes, followed by 65°C for 30 minutes, to polish the ends of the chromatin fragments. Bridge Ligation Mix and Bridge Ligase (Cat # PN DG-NUC-001, Dovetail Genomics) were subsequently added, and the samples were incubated at 22°C for 30 minutes to initiate ligation. Intra-aggregate Ligation Buffer and Enzyme Mix (Cat # PN DG-NUC-001, Dovetail Genomics) were added, with incubation at 22°C for an additional hour to facilitate intra-aggregate ligation. To isolate DNA, reverse cross-linking was performed using Proteinase K and Crosslink Reversal Buffer. The samples were incubated at 55°C for 15 minutes and 68°C for 45 minutes. Purified DNA was then prepared for library construction. End Repair Master Mix (Cat # PN DG-LIB-001, Dovetail Genomics) was added to the DNA and incubated at 20°C for 30 minutes, followed by 65°C for 30 minutes for end repair. For adapter ligation, Illumina-compatible adapters were added along with ligation enzyme mix and enhancer, and the DNA was purified using SPRIselect beads (Cat # B23317, Beckman Coulter, USA). Adaptor-ligated DNA was washed and captured with Streptavidin beads (Cat # PN DG-REF-001, Dovetail Genomics). Following capture, PCR amplification was performed, and fragments within a 350–1,000 bp size range were selected for final library construction. Prepared libraries were sequenced on an Illumina sequencing platform.

## Data availability

The datasets generated during the current study are available in the GEO repository under accession number, GSE291291.

## Code availability

Our method is implemented as a python package and is freely available at Under GNU General Public License (GPL-v3.0) (https://github.com/KunFang93/Nuc3DMap).

## Authors’ contributions

VXJ conceived the project. KF developed the algorithm and tool, Nuc3DMap. TL assisted in the data analyses. LC conducted the Micro-C experiment. PAK provided the assistance in implementing the ensembled LOESS-sparse Knight-Ruiz algorithm. VXJ, KF, TL and LC wrote the manuscript, with all authors contributing to writing and providing the feedback. All authors read and approved the final manuscript.

## Acknowledgements

We thank the UTHSA Next Generation Sequencing Facilities, Dr. Zhao Lai for producing the Micro-C data. We would also thank Drs. Nils Krietenstein and Nezar Abdennur and Open2C for the helpful clarification and suggestions.

