## Supplementary material for "Nuc3DMap delineates finer nucleosome-based three-dimensional chromatin architecture": Suppl. Materials

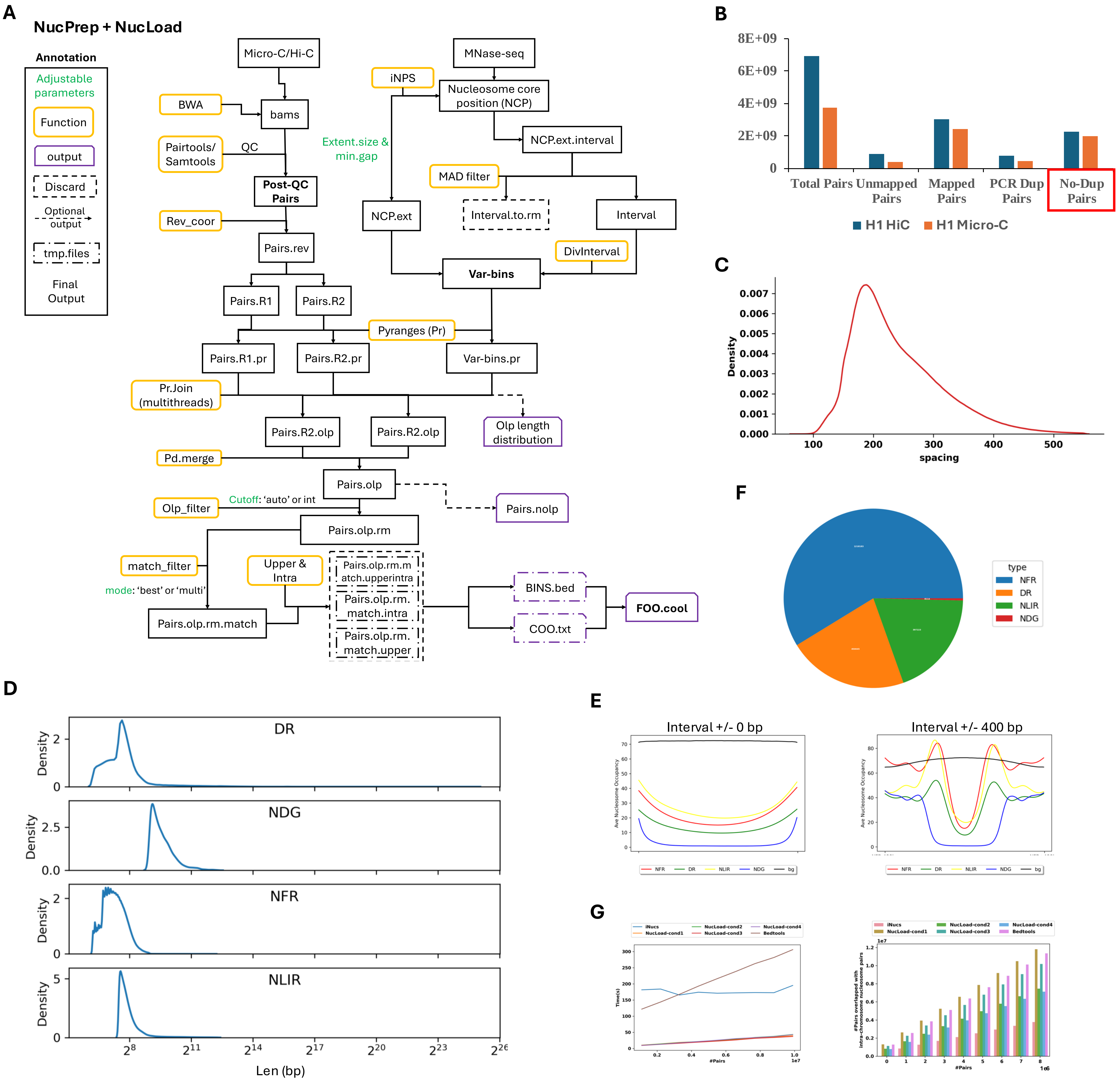
A
NucPrep + NucLoad
Annotation
Adjustable parameters
Function
output
Discard
Optional output
tmp.files
Final Output
Micro-C/Hi-C
BWA
bams
Pairtools/ Samtools
QC
Post-QC Pairs
Rev_coor
Pairs.rev
Pairs.R1
Pairs.R2
MNase-seq
iNPS
Nucleosome core position (NCP)
NCP.ext.interval
Extent.size & min.gap
MAD filter
NCP.ext
Interval.to.rm
Interval
DivInterval
Var-bins
Pyranges (Pr)
Pairs.R1.pr
Pairs.R2.pr
Var-bins.pr
Pr.Join (multithreads)
Pairs.R2.olp
Pairs.R2.olp
Olp length distribution
Pd.merge
Pairs.olp
Pairs.nolp
Olp_filter
Cutoff: 'auto' or int
Pairs.olp.rm
match_filter
mode: 'best' or 'multi'
Upper & Intra
Pairs.olp.rm.match
Pairs.olp.rm.match.upperintra
Pairs.olp.rm. match.intra
Pairs.olp.rm. match.upper
BINS.bed
COO.txt
FOO.cool
B
8E+09
6E+09
4E+09
2E+09
0
Total Pairs
Unmapped Pairs
Mapped Pairs
PCR Dup Pairs
No-Dup Pairs
H1 HiC
H1 Micro-C
C
Density
spacing
F
type
NFR
DR
NLIR
NDG
D
DR
NDG
NFR
NLIR
Density
Len (bp)
E
Interval +/- 0 bp
Interval +/- 400 bp
Ave Nucleosome Occupancy
NFR
DR
NLIR
NDG
bg
G
iNucs
NucLoad-cond1
NucLoad-cond2
NucLoad-cond3
NucLoad-cond4
Bedtools
Time(s)
#Pairs
#Pairs overlapped with intra-chromosome nucleosome pairs

### Extended Data Fig. 1.

**A.** The detailed workflow of NucPrep and NucLoad.

**B.** The bar plot showing the number of total pairs, unmapped pairs, mapped pairs, PCR duplicated pairs, and no-duplicated pairs. No-duplicated is the final output for NucPrep.

**C.** The size distribution of the spacing of N-bins.

**D.** The size distribution of the four types of Is.

**E.** The MNase-seq signals of four types of Is with 0 and 400 bp extension

**F.** The pie chart showing the proportion of four types of Is.

**G.** The line plot showing the speed comparison among, iNucs, Bedtools, NucLoad-cond1 (olp_cutoff =1, match_mode: multi), NucLoad_cond2 (olp_cutoff =1, match_mode: best), NucLoad_cond3 (olp_cutoff = auto, match_mode: multi) and NucLoad_cond4 (olp_cutoff = auto, match_mode: best). The bar plot demonstrates the accuracy comparison among iNucs, Bedtools, NucLoad-cond1 (olp_cutoff =1, match_mode: multi), NucLoad_cond2 (olp_cutoff =1, match_mode: best), NucLoad_cond3 (olp_cutoff = auto, match_mode: multi) and NucLoad_cond4 (olp_cutoff = auto, match_mode: best).

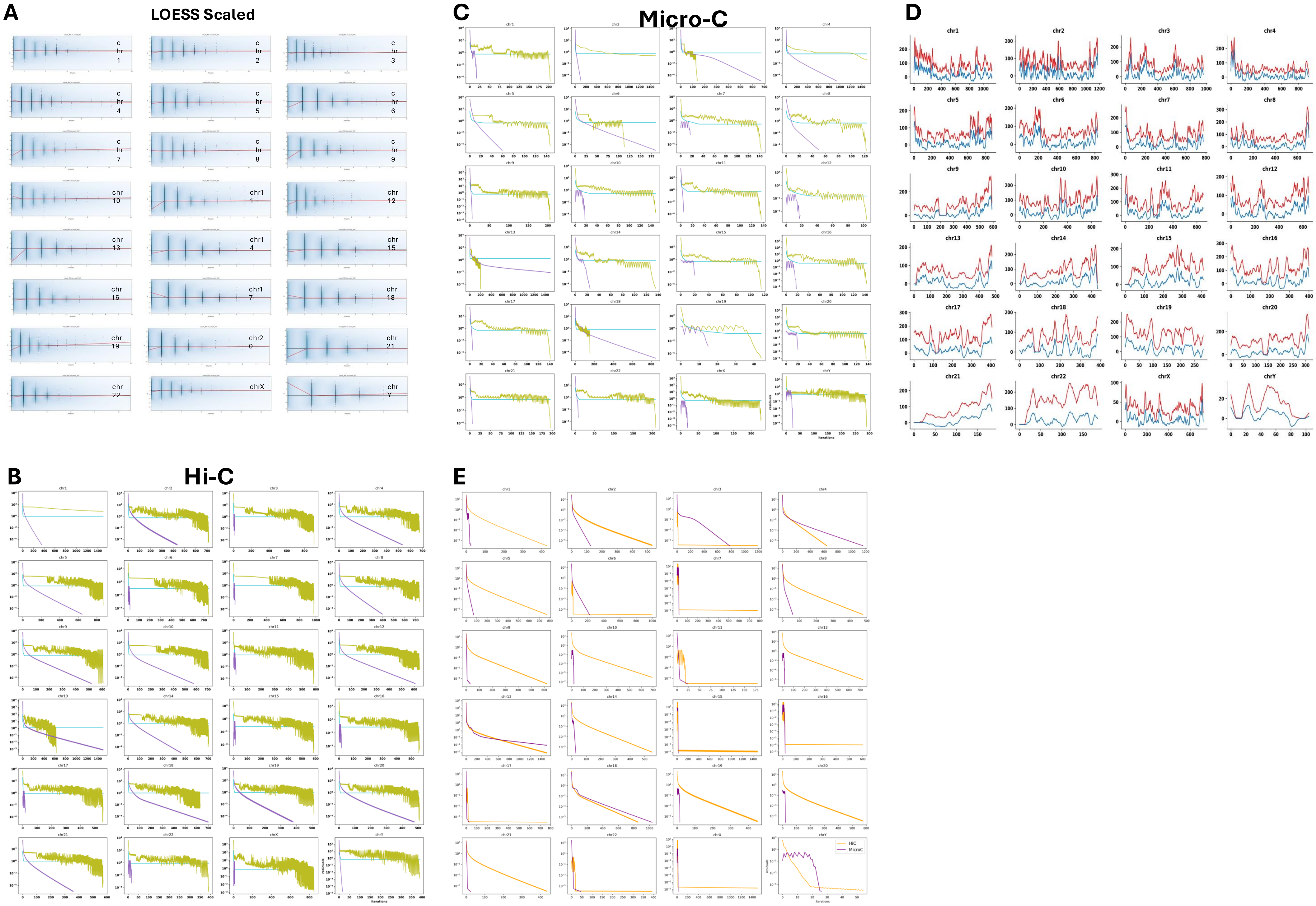
A
LOESS Scaled
C
Micro-C
D
B
Hi-C
E
HiC
MicroC

### Extended Data Fig. 2.

**A.** M-DS plots of each chromosome showing a successful scaling after DS-Anchor scaling step.

**B.** Line plots showing the convergency of three matrix balancing methods (RAS, KR and sKR) on Hi-C contact matrix.

**C.** Line plots showing the convergency of three matrix balancing methods (RAS, KR and sKR) on Micro-C contact matrix.

**D.** Balanced Hi-C and Micro-C matrices with scaled marginal values by sKR.

**E.** The marginal values of each chromosome before and after the normalization of NucMerge.

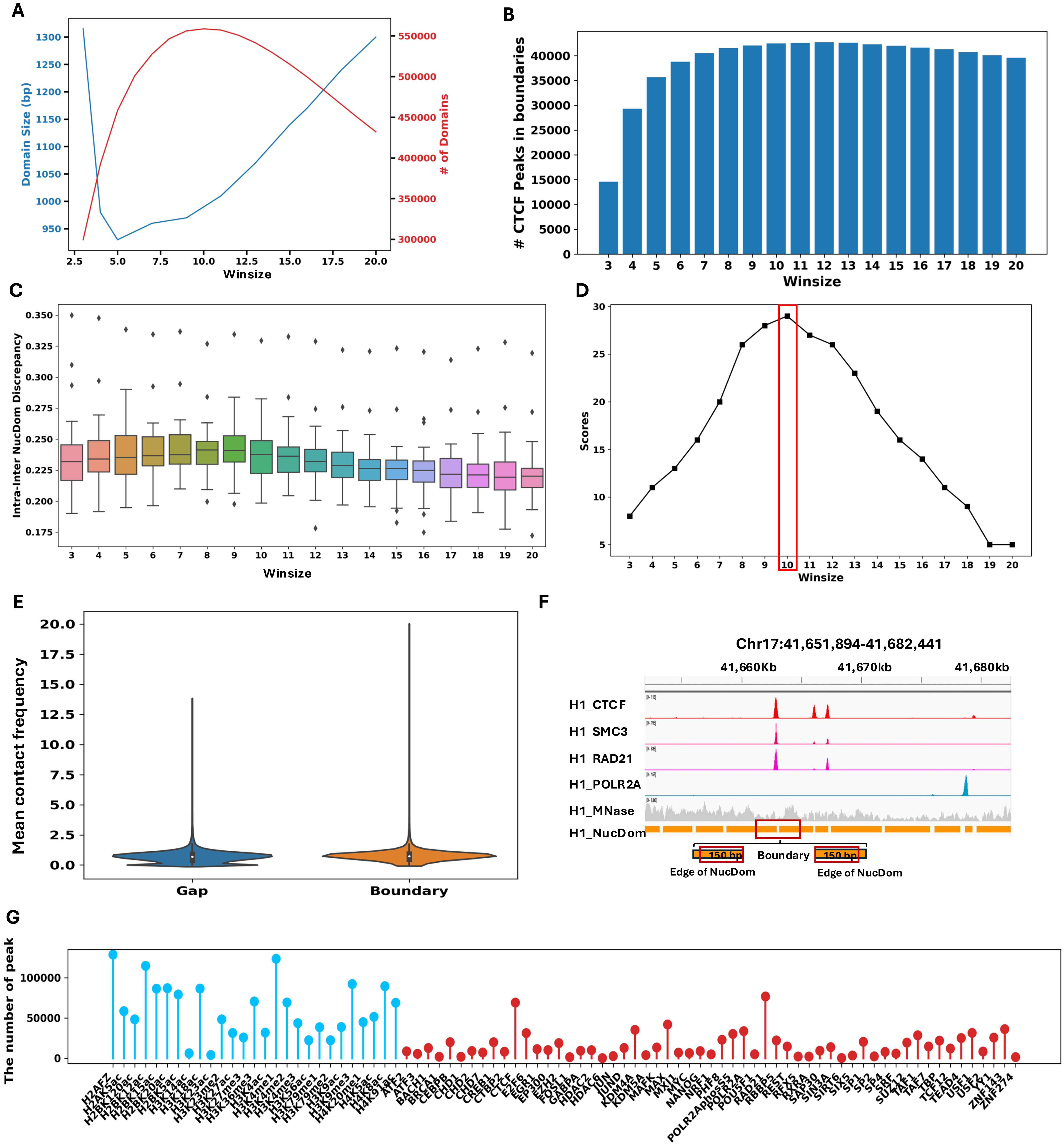

A
Domain Size (bp)
### of Domains
Winsize
B
### CTCF Peaks in boundaries
Winsize
C
Intra-Inter NucDom Discrepancy
Winsize
D
Scores
Winsize
E
Mean contact frequency
Gap
Boundary
F
Chr17:41,651,894-41,682,441
41,660Kb
41,670kb
41,680kb
H1_CTCF
H1_SMC3
H1_RAD21
H1_POLR2A
H1_MNase
H1_NucDom
150 bp
Boundary
150 bp
Edge of NucDom
Edge of NucDom
G
The number of peak

### Extended Data Fig. 3.

**A.** The double-axis line plot showing the size of NucDs and the number of NucBs under different Winsize.

**B.** The bar plot showing the number of CTCF peaks located on the NucBs identified under different Winsize.

**C.** The box plot showing the inter-intra contact frequency discrepancy of NucDs under different Winsize.

**D.** The line plot showing the Winsize of 10 has the highest score, suggesting it is the optimal parameter.

**E**. The violin plot showing the mean contact frequency of NucG and NucB.

**F.** The IGV screenshot showing NucB and the definition of the edge of NucD.

**G.** The lollipop-plot showing the number of peaks in each of curated histone modifications and TFs.

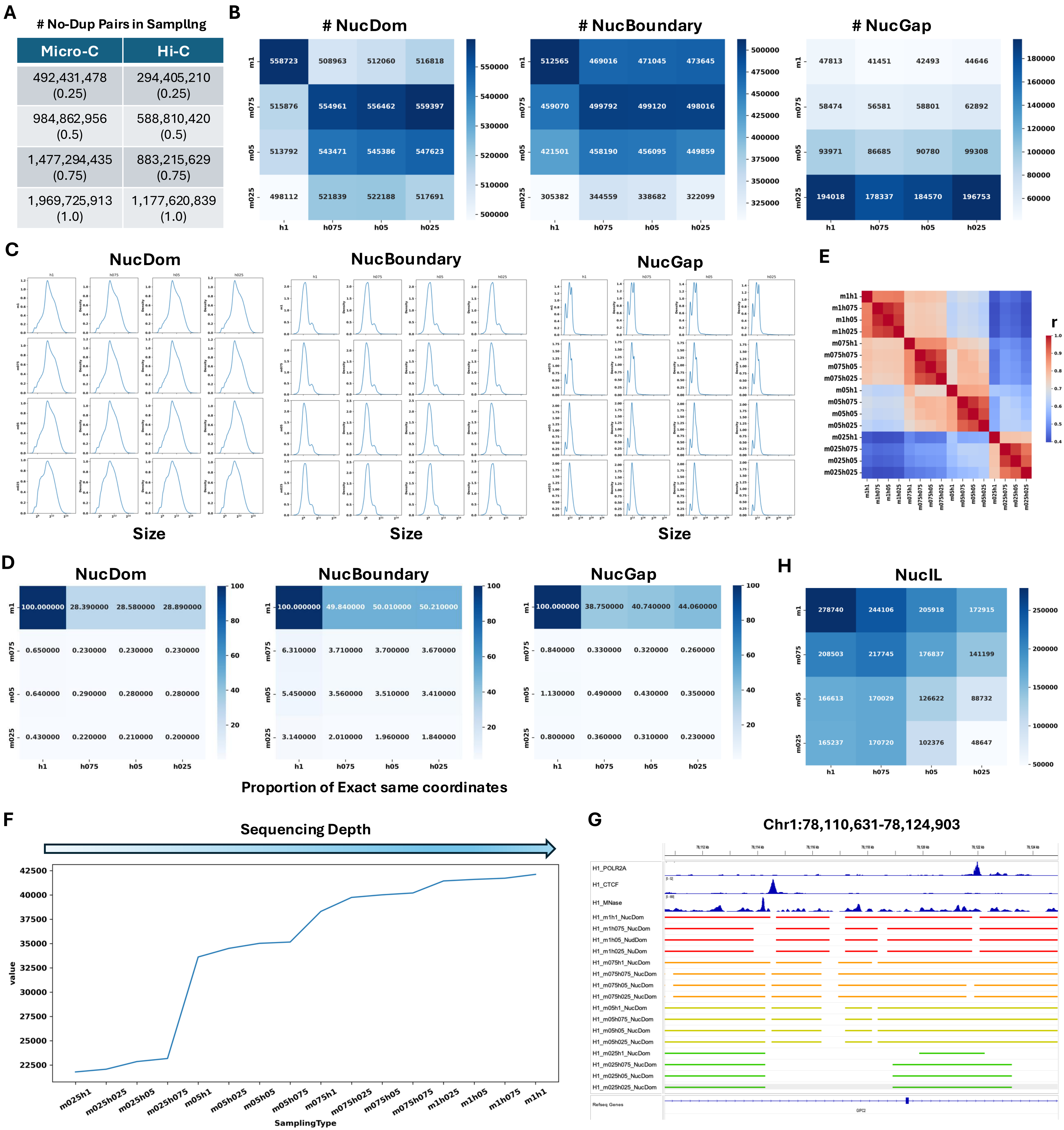
A
### No-Dup Pairs in Sampling
Micro-C
Hi-C
492,431,478 (0.25)
294,405,210 (0.25)
984,862,956 (0.5)
588,810,420 (0.5)
1,477,294,435 (0.75)
883,215,629 (0.75)
1,969,725,913 (1.0)
1,177,620,839 (1.0)
B
### NucDom
558723 508963 512060 516818
515876 554961 556462 559397
513792 543471 545386 547623
498112 521839 522188 517691
### NucBoundary
512565 469016 471045 473645
459070 499792 499120 498016
421501 458190 456095 449859
305382 344559 338682 322099
### NucGap
47813 41451 42493 44646
58474 56581 58801 62892
93971 86685 90780 99308
194018 178337 184570 196753
m1
m075
m05
m025
h1
h075
h05
h025
C
NucDom
NucBoundary
NucGap
Size
E
r
D
NucDom
100.000000 28.390000 28.580000 28.890000
0.650000 0.230000 0.230000 0.230000
0.640000 0.290000 0.280000 0.280000
0.430000 0.220000 0.210000 0.200000
NucBoundary
100.000000 49.840000 50.010000 50.210000
6.310000 3.710000 3.700000 3.670000
5.450000 3.560000 3.510000 3.410000
3.140000 2.010000 1.960000 1.840000
NucGap
100.000000 38.750000 40.740000 44.060000
0.840000 0.330000 0.320000 0.260000
1.130000 0.490000 0.430000 0.350000
0.800000 0.360000 0.310000 0.230000
Proportion of Exact same coordinates
H
NucIL
278740 244106 205918 172915
208503 217745 176837 141199
166613 170029 126622 88732
165237 170720 102376 48647
F
Sequencing Depth
value
SamplingType
m025h1
m025h025
m025h05
m025h075
m05h1
m05h025
m05h05
m05h075
m075h1
m075h025
m075h05
m075h075
m1h025
m1h05
m1h075
m1h1
G
Chr1:78,110,631-78,124,903
H1_POLR2A
H1_CTCF
H1_MNase
H1_m1h1_NucDom
H1_m1h075_NucDom
H1_m1h05_NudDom
H1_m1h025_NuDom
H1_m075h1_NucDom
H1_m075h075_NucDom
H1_m075h05_NucDom
H1_m075h025_NucDom
H1_m05h1_NucDom
H1_m05h075_NucDom
H1_m05h05_NucDom
H1_m05h025_NucDom
H1_m025h1_NucDom
H1_m025h075_NucDom
H1_m025h05_NucDom
H1_m025h025_NucDom
Refseq Genes

#### Extended Data Fig. 4.

- **A**. The table presenting the number of reads after down-sampling for both Hi-C and Micro-C, with four tiers of down-sampling applied to each: 0.25, 0.5, 0.75, and 1.0.
- **B**. Heatmaps showing the impact of down-sampling on the number of identified NucD, NucB, and NucG.
- **C**. Line plots showing the effect of down-sampling on the size of identified NucD, NucB and NucG.
- **D**. Heatmaps showing the impact of down-sampling on the percentage of overlapped NucD, NucB, and NucG compared the Micro-C 1.0 and Hi-C 1.0.
- **E**. The heatmap displaying the Pearson correlation coefficients among 16 different combinations of Micro-C and Hi-C down-sampling levels, highlighting the depth of Micro-C dominant the NucDom performance.
- **F**. The line plot illustrates the number of CTCF sites captured within identified NucB across varying sequencing depth conditions, suggesting the low depth (25%) impair the performance of the NucDom significantly.
- **G**. The IGV screenshot visualizing the NucD and NucB detected within Chr1:78,110,631-78,124,903.
- **H**. The heatmap showing the impact of down-sampling on the number of NucIL.

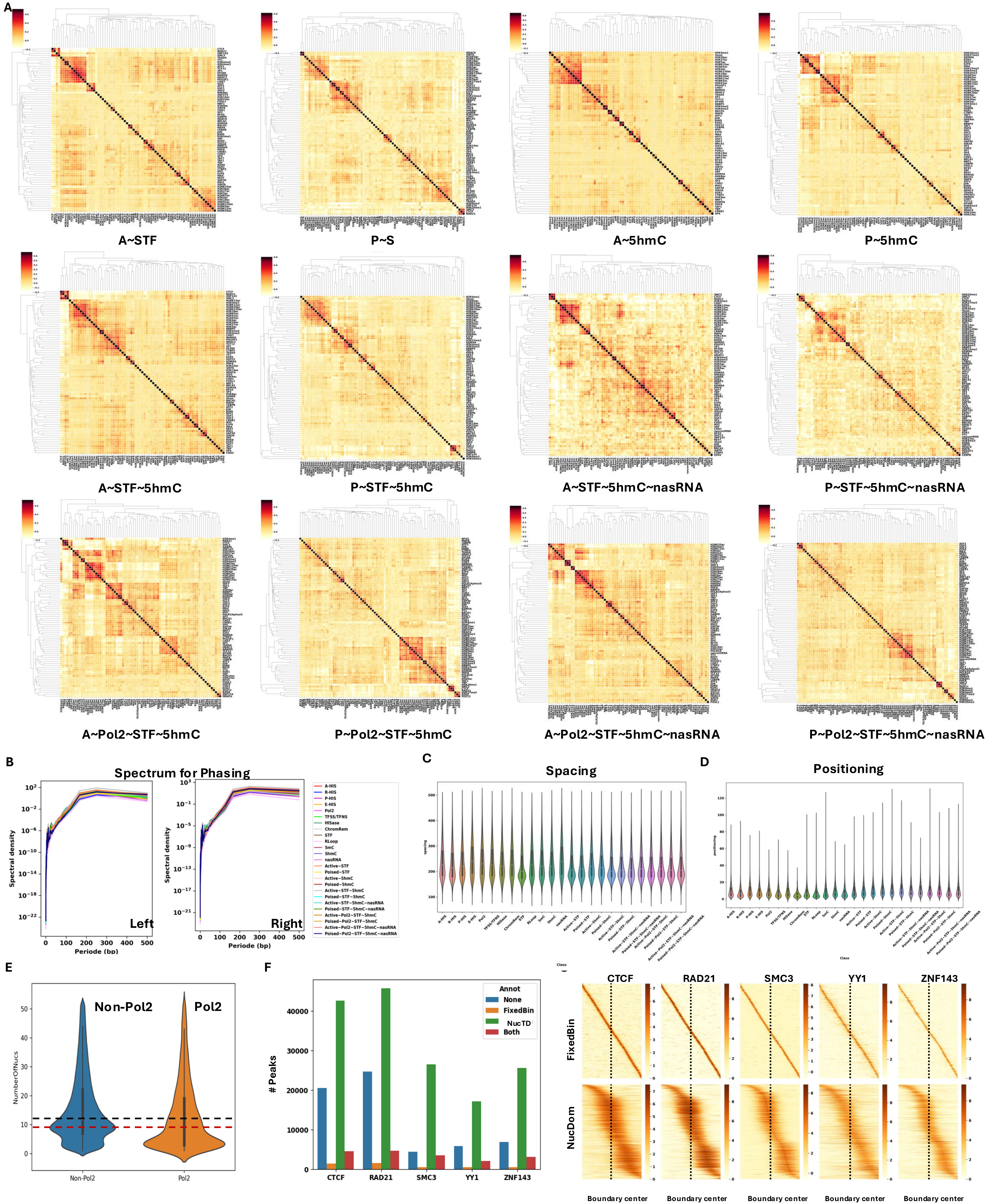
A
A~STF
P~S
A~5hmC
P~5hmC
A~STF~5hmC
P~STF~5hmC
A~STF~5hmC~nasRNA
P~STF~5hmC~nasRNA
A~Pol2~STF~5hmC
P~Pol2~STF~5hmC
A~Pol2~STF~5hmC~nasRNA
P~Pol2~STF~5hmC~nasRNA
B
Spectrum for Phasing
Spectral density
Periode (bp)
Left
Right
A-HIS
R-HIS
P-HIS
E-HIS
Pol2
TFSS/TFNS
HISase
ChromRem
STF
RLoop
5mC
5hmC
nasRNA
Active~STF
Poised~STF
Active~5hmC
Poised~5hmC
Active~STF~5hmC
Poised~STF~5hmC
Active~STF~5hmC~nasRNA
Poised~STF~5hmC~nasRNA
Active~Pol2~STF~5hmC
Poised~Pol2~STF~5hmC
Active~Pol2~STF~5hmC~nasRNA
Poised~Pol2~STF~5hmC~nasRNA
C
Spacing
D
Positioning
E
Non-Pol2
Pol2
NumberOfNucs
F
### Peaks
Annot
None
FixedBin
NucTD
Both
CTCF
RAD21
SMC3
YY1
ZNF143
FixedBin
NucDom
Boundary center

### Extended Data Fig. 5.

**A.** The correlation matrices of different combination types of NucB.

**B.** The spectrum density plots for each type of NucB.

**C.** The violin plots showing the distribution of the nucleosome spacing score of each type of NucB.

**D.** The violin plots showing the distribution of the nucleosome positioning score of each type of NucB.

**E.** The violin plots showing the distribution of the number of nucleosomes associated with Pol2 and non-Pol2 NucDs.

**F.** The bar plot (left) showing the number of CTCFs located at NucBs and Boundaries curated from 4DN, respectively. The density plot (right) shows CTCFs were more concentrated at NucBs.

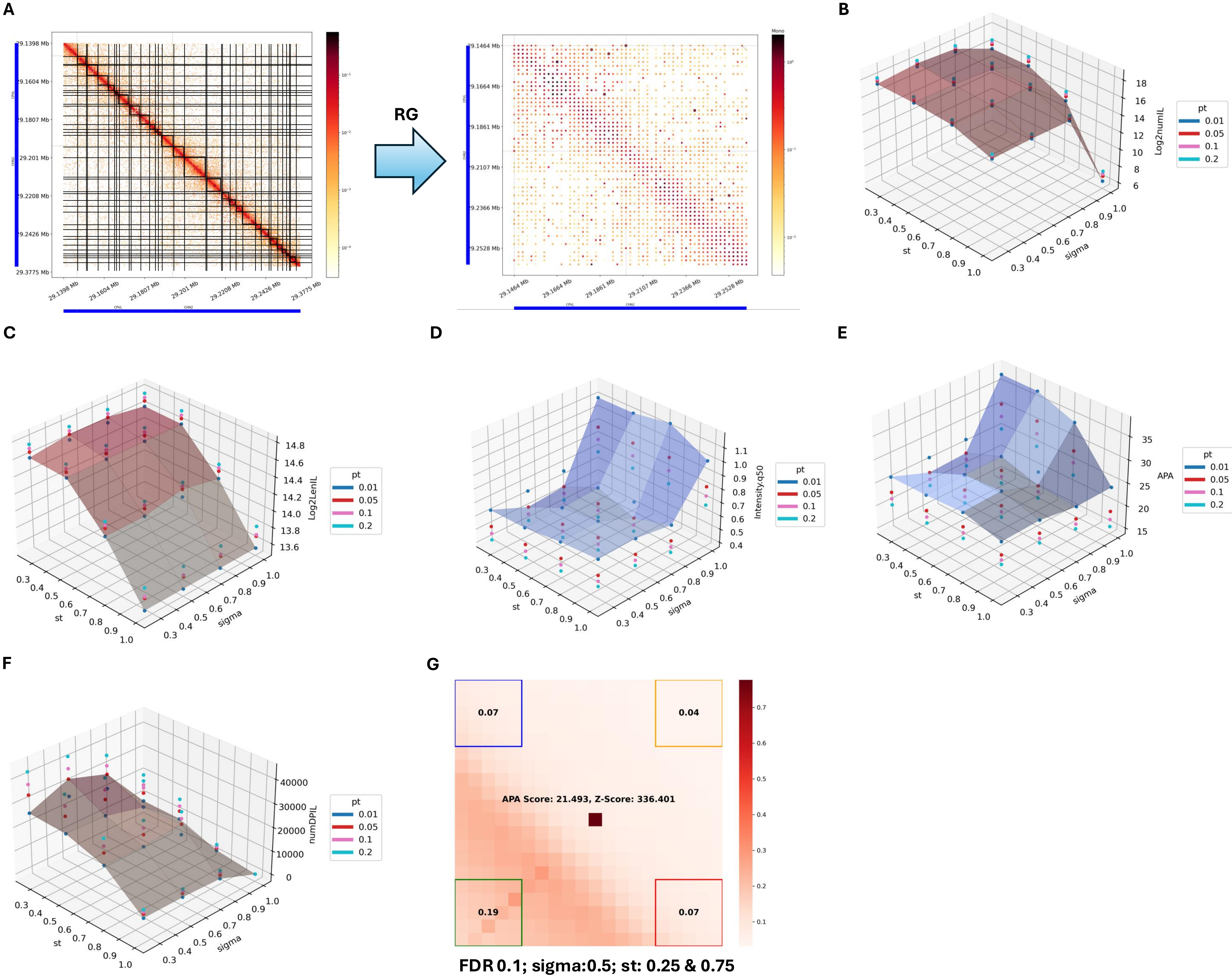
A
B
C
D
E
F
G
RG
29.1398 Mb
29.1604 Mb
29.1807 Mb
29.201 Mb
29.2208 Mb
29.2426 Mb
29.3775 Mb
29.1464 Mb
29.1664 Mb
29.1861 Mb
29.2107 Mb
29.2366 Mb
29.2528 Mb
Mono
Log2numIL
Log2LenIL
Intensity.q50
APA
numDPIL
st
sigma
pt
0.01
0.05
0.1
0.2
0.07
0.04
0.19
0.07
APA Score: 21.493, Z-Score: 336.401
FDR 0.1; sigma:0.5; st: 0.25 & 0.75

### Extended Data Fig. 6.

**A.** A nucleosome-based contact matrix being converted into a re-normalization grouping (RG) matrix with NucD/NucB/NucG identified in NucDom.

**B.** The number of NucILs identified under different combinations of st (spatial threshold), sigma0 (initial sigma for SIFT), and pt (pval threshold). A total of 64 combinations were tested, including four st values (0.25, 0.5, 0.75, and 1), four pt values (0.01, 0.05, 0.1, and 0.2), as well as four sigma0 values (0.25, 0.5, 0.75 and 1.0).

**C.** The length of NucIL identified under different combinations of st, sigma0, and pt.

**D.** The median intensity of NucIL identified under different combinations of st, sigma0, and pt.

**E.** The APA (aggregated peak analysis) score of NucIL identified under different combinations of st, sigma0, and pt.

**F.** The number of D-P NucL identified under different combinations of st, sigma0, and pt.

**G.** The APA score of NucIL with the optimal parameter.

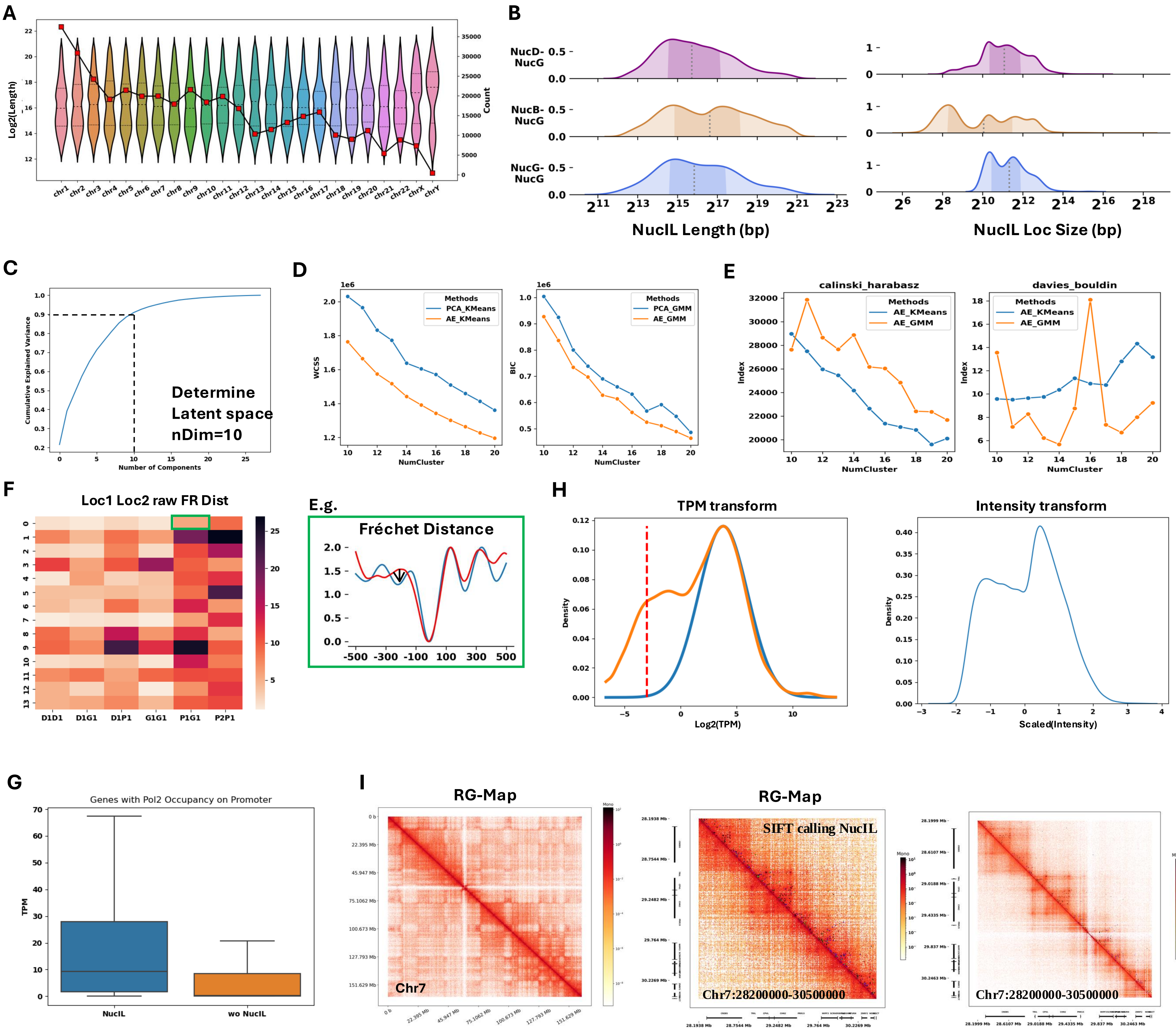
A
Log2(Length)
Count
B
NucD-NucG
NucB-NucG
NucG-NucG
NucIL Length (bp)
NucIL Loc Size (bp)
C
Cumulative Explained Variance
Number of Components
Determine
Latent space
nDim=10
D
WCSS
BIC
NumCluster
Methods
PCA_KMeans
AE_KMeans
PCA_GMM
AE_GMM
E
calinski_harabasz
davies_bouldin
Index
NumCluster
AE_KMeans
AE_GMM
F
Loc1 Loc2 raw FR Dist
D1D1
D1G1
D1P1
G1G1
P1G1
P2P1
E.g.
Fréchet Distance
H
TPM transform
Intensity transform
Density
Log2(TPM)
Scaled(Intensity)
G
Genes with Pol2 Occupancy on Promoter
TPM
NucIL
wo NucIL
I
RG-Map
Chr7
SIFT calling NucIL
Chr7:28200000-30500000

### Extended Data Fig. 7.

**A.** A double-axis plot showing the length distribution of NucIL (violin plot) alongside the count of NucIL (line plot) across each chromosome.

**B.** The ridge plot displaying the length distribution (left) and loci size distribution (right) for NucD-NucG, NucB-NucG, and NucG-NucG types of NucILs.

**C.** The dimension of the latent space determined by cumulative explained variance.

**D.** Within-cluster sum-of-squares (WCSS) and Bayesian Information Criterion (BIC) suggesting that the autoencoder (AE) outperforms PCA when combined with either clustering method (KMeans or GMM). Our evaluation indicated superior performance of the AE configurations, with AE-KMeans displaying consistently lower WCSS and AE-GMM showing lower BIC values across all tested cluster counts.

**E.** Calinski-Harabasz and Davies–Bouldin indices indicating that AE combined with GMM outperforms AE + KMeans for most cluster numbers, suggesting that 14 is the optimal number of clusters.

**F.** The heatmap (left) showing the raw Fréchet distance (FR) values for each cluster, categorized by genomic location. The enrichment plot (right) illustrates the process of calculating FR values.

**G.** The boxplot displaying the distribution of TPM values between genes associated with PolII-related NucLs and genes associated with PolII without NucLs.

**H.** Line plots showing the distribution of scaled TPM and NucL intensity using zTPM and Box-Cox transformations.

**I.** Visualization of the RGMap for Chr7, a section of Chr7 (Chr7: 28,200,000–30,500,000), and the corresponding nucleosome-based map for the same region, with NucILs labeled.

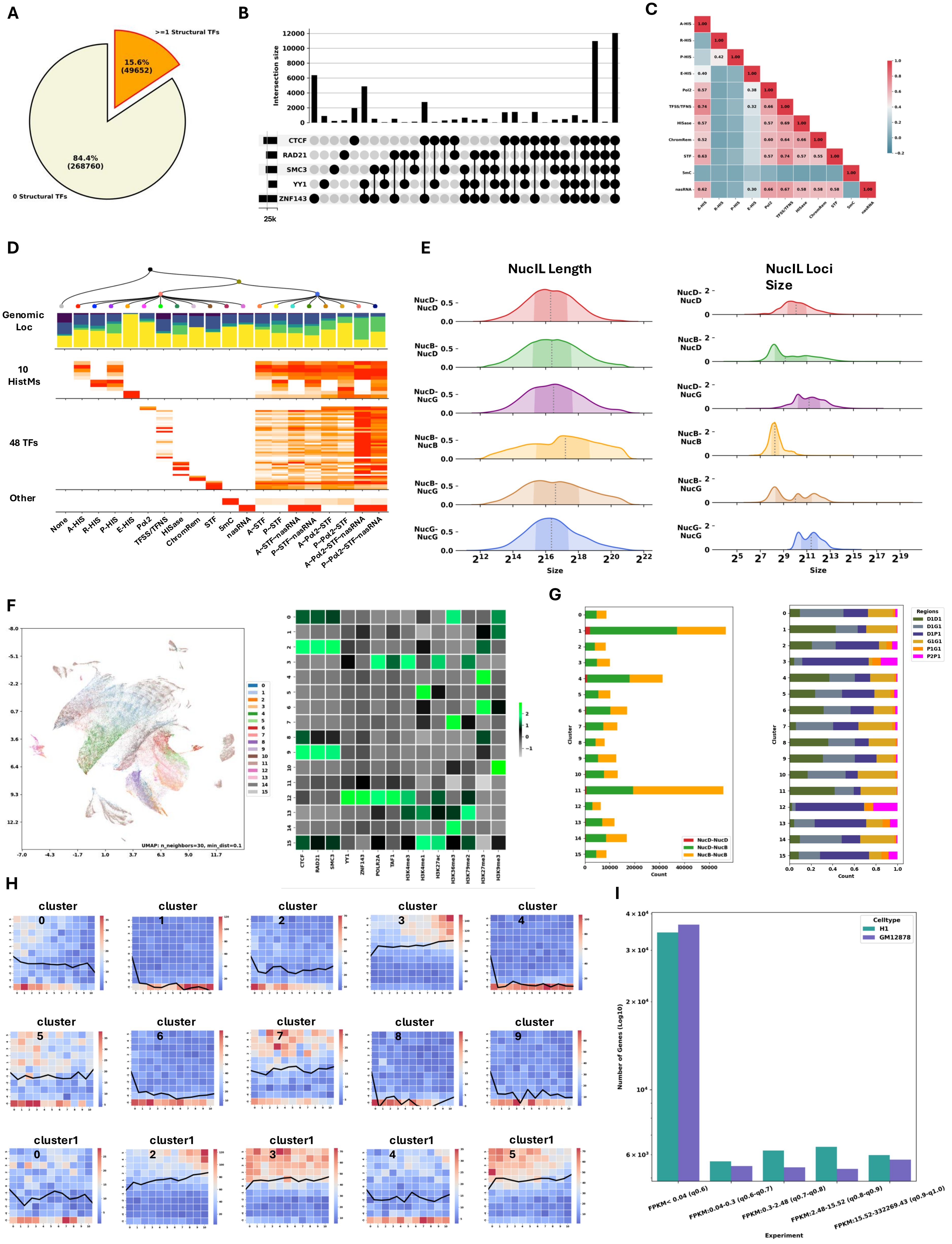

A
>=1 Structural TFs
15.6% (49652)
84.4% (268760)
0 Structural TFs
B
Intersection size
CTCF
RAD21
SMC3
YY1
ZNF143
25k
C
D
Genomic Loc
10 HistMs
48 TFs
Other
E
NucIL Length
NucIL Loci Size
Size
F
UMAP: n_neighbors=30, min_dist=0.1
G
Cluster
Count
NucD-NucD
NucD-NucB
NucB-NucB
Regions
D1D1
D1G1
D1P1
G1G1
P1G1
P2P1
H
I
Celltype
H1
GM12878
Number of Genes (Log10)
Experiment

### Extended Data Fig. 8.

**A.** Pie chart showing the proportion of NucBs associated with structural-related transcription factors (STFs) in GM12878 cells.

**B.** UpSet plot displaying combinations of STF within STF-related NucBs in GM12878 cells.

**C.** The heatmap representing the correlation metrics across 11 ‘single’ type of NucBs in GM12878 cells.

**D.** Multi-feature plot detailing genomic characteristics of 19 different NucBs types (11 single, 9 combination), including genomic location (desert, distal, proximal, promoter and genebody), associated histone modifications, TFs, 5mC and nasRNA .

**E.** The KDE-plot (left) revealing the length distribution of six NucIL types: NucD-NucD, NucD-NucB, NucB-NucB, NucD-NucB, NucD-NucG, NucB-NucG, and NucG-NucG. The plot (right) panel shows the size distribution of these types.

**F.** The UMAP (left) visualizing clusters of NucILs that were conducted using an AE and a GMM for clustering, ased on selected histone modifications and TFs at their interacting loci. The heatmap (right) displays the centroids of each cluster, highlighting distinct combinatorial patterns of histone modifications and TFs.

**G.** The bar plot (left) elucidating the composition of NucIL clusters, segmented by interaction types: NucD-NucD, NucD-NucB, and NucB-NucB. The stack plot (right) re-categorizes NucIL into NucL by the genomic location of their interacting loci, including D1D1, D1P1, D1G1, P1G1, G1G1, and P2P1.

**H.** The expression-interaction matrix for different NucIL clusters in GM12878 cells.

**I.** The number of genes stratified by FPKM levels in H1 and GM12878 cells, respectively.
